# Evidence for widespread selection in shaping the genomic landscape during speciation of *Populus*

**DOI:** 10.1101/819219

**Authors:** Jing Wang, Nathaniel R. Street, Eung-Jun Park, Jianquan Liu, Pär K. Ingvarsson

## Abstract

Increasing our understanding of how various evolutionary processes drive the genomic landscape of variation is fundamental to a better understanding of the genomic consequences of speciation. However, the genome-wide patterns of within- and between-species variation have not been fully investigated in most forest tree species despite their global ecological and economic importance. Here, we use whole-genome resequencing data from four *Populus* species spanning the speciation continuum to reconstruct their demographic histories, investigate patterns of diversity and divergence, infer their genealogical relationships and estimate the extent of ancient introgression across the genome. Our results show substantial variation in these patterns along the genomes although this variation is not randomly distributed but is strongly predicted by the local recombination rates and the density of functional elements. This implies that the interaction between recurrent selection and intrinsic genomic features has dramatically sculpted the genomic landscape over long periods of time. In addition, our findings provide evidence that, apart from background selection, recent positive selection and long-term balancing selection are also crucial components in shaping patterns of genome-wide variation during the speciation process.

## Introduction

Determining the evolutionary forces affecting patterns of genome-wide variation has been a central goal in evolutionary biology over the past several decades (Seehausen et al., 2014). Furthermore, studying variation in levels of differentiation within and between closely related species has the potential to yield important insights into the process of speciation (Ravinet et al., 2017; Wolf & Ellegren, 2017). Studies in a broad range of taxonomic groups have revealed a picture of a highly heterogeneous genomic landscape with peaks and valleys of diversity and differentiation (Han et al., 2017; Nadeau et al., 2012; Stankowski et al., 2019; Turner, Hahn, & Nuzhdin, 2005). Local peaks of elevated divergence are usually referred to as ‘speciation islands’ and are thought to represent regions that drive the reproductive isolation between incipient species (Abbott et al., 2013; Wu, 2001). Between these islands, gene flow acts to homogenize the reminder of genome and hence acts to limit differentiation (Feder, Egan, & Nosil, 2012; Nosil, Funk, & Ortiz - Barrientos, 2009). However, a plethora of recent studies highlight that the heterogeneous patterns of differentiation can evolve through processes that are unrelated to speciation *per se* (Burri et al., 2015; Cruickshank & Hahn, 2014). For example, even in the absence of gene flow, natural selection, in the form of either a selective sweep or background selection, can cause reduced genetic diversity not only at the target sites under selection but also at linked neutral sites (Han et al., 2017; Phung, Huber, & Lohmueller, 2016). Such selection could accelerate lineage sorting and will hence inevitably result in increased genetic differentiation between species in these regions (Burri, 2017). Furthermore, the long-term action of linked selection in ancestral as opposed to extant lineages can also affect the amount and distribution of ancestral polymorphisms (Ma et al., 2018; Munch, Nam, Schierup, & Mailund, 2016; Scally et al., 2012), which can further result in heterogeneous patterns of genealogical relationships among closely related species (Mailund, Munch, & Schierup, 2014; Pease & Hahn, 2013). Despite widespread interest in speciation genomics, there remains little consensus as to how various evolutionary processes have shaped the genomic landscape during the speciation process that eventually gives rise to new species (Ravinet et al., 2017).

Empirical studies suggest that the formation of the genomic landscape of diversity during speciation is highly influenced by the demographic histories of the species, the types of selection acting on different genomic regions and also several other intrinsic genomic features (Burri, 2017; Ellegren & Galtier, 2016). Disentangling the effects of speciation (i.e. species split time, strict isolation or divergence with gene flow) is important for interpreting the patterns of genome-wide variation, because without a clear picture of the demographic history of the descendant species, it is challenging to distinguish whether heterogenous genomic differentiation arose due to genetic drift, local adaptation or introgression (Nadachowska-Brzyska et al., 2013; Ravinet et al., 2018). Furthermore, as the speciation process advances, the evolution of genome-wide patterns of variation can be influenced by different forms of selection (Cutter & Payseur, 2013). Under a background selection model, purifying selection continuously eliminates deleterious mutations, resulting in reduced levels of genetic diversity at linked loci and increased levels of *F*_ST_ (a relative measure of genetic divergence) (Charlesworth, 2012; Charlesworth, Morgan, & Charlesworth, 1993; Hudson & Kaplan, 1995). Under a selective sweep model, genetic variants linked to beneficial mutations acted upon by positive selection hitchhike along and reach high frequency (Kaplan, Hudson, & Langley, 1989; Smith & Haigh, 1974). Accordingly, even in the absence of gene flow, selection due to, for instance local ecological adaptation, can result in reduced diversity and increased *F*_ST_ (Cruickshank & Hahn, 2014). In comparison to purifying and positive selection, long-term balancing selection favors the maintenance of advantageous polymorphisms for many generations, which instead result in genomic regions with elevated genetic diversity and reduced *F*_ST_ (Charlesworth, 2006; Guerrero & Hahn, 2017). As deleterious mutations are assumed to be much more common compared to beneficial mutations, background selection has been argued to play a major role in the evolution of diversity (Burri, 2017; Lohmueller et al., 2011; Phung et al., 2016). However, many recent simulation studies have shown that background selection alone is far from sufficient for generating the heterogenous genomic landscapes observed in empirical studies of recently diverged species, and other evolutionary processes (such as positive selection) are thus required to explain the observed patterns (Matthey - Doret & Whitlock 2019; Stankowski et al., 2019).

Regardless of the role of demographic processes and selection, genomic features, such as recombination rate variation and the heterogeneous density of functional sites, are also expected to play key roles in mediating the efficacy and extent of selection and gene flow, as well as how these processes interact as the speciation process proceeds (Flaxman, Wacholder, Feder, & Nosil, 2014; Hurst, Pál, & Lercher, 2004; Nachman & Payseur, 2012). Local rates of recombination interacts with natural selection and are known to have a profound effect on patterns of genomic diversity, incomplete lineage sorting (ILS) and rates of introgression (Begun & Aquadro, 1992; Comeron, Williford, & Kliman, 2008; Cutter & Payseur, 2013). Independent of the recombination rate, the density of functional sites can also influence genome-wide patterns of diversity since functional regions are more likely to experience either stronger effects of positive or purifying selection compared to nonfunctional regions where mutations are assumed to have little effect on fitness (Al-Shahrour et al., 2010; Nordborg et al., 2005). The long-term diversity-reducing effects of selection in functional regions will reduce locally effective population size (*N*_e_), accelerate lineage sorting and increase genetic divergence between species (Flowers et al., 2011; Hobolth, Dutheil, Hawks, Schierup, & Mailund, 2011). As it becomes increasingly feasible to generate whole genome resequencing data from closely related species, the importance of conserved genomic features in shaping the topography of the genomic landscape of speciation has increasingly been highlighted by several studies in a diverse set of taxa showing highly correlated patterns of differentiation among independently species pairs (Burri, 2017; Delmore et al., 2018; Van Doren et al., 2017; Vijay et al., 2017).

Forest trees provide an excellent system to address the genomic architecture of adaptation and speciation in natural populations because they are mostly undomesticated without much anthropogenic influence, ecologically important across a wide variety of habitats and harbour abundant genetic and phenotypic variation (Neale & Ingvarsson, 2008; Neale & Kremer, 2011). In this study, we focus on four *Populus* species (*Populus tremula*, *P. davidiana*, *P. tremuloides* and *P. trichocarpa*) that span the speciation continuum. All four species are all deciduous, obligated outcrossing tree species that have wide geographical distributions throughout the Northern Hemisphere (Figure 1A). Among them, *P. tremula* (European aspen), *P. davidiana* (Chinese aspen) and *P. tremuloides* (American aspen) are sibling aspen species belonging to the same section of the genus *Populus* (section *Populus*) (Eckenwalder, 1996; Hamzeh & Dayanandan, 2004). Earlier phylogenetic studies have revealed that *P. tremuloides* diverged from the other two species following the break-up of the Bering Land bridge, whereas the uplift of the Qinghai-Tibetan Plateau and the associated climate oscillations may have driven the divergence between *P. tremula* and *P. davidiana* (Du et al., 2015). In addition, these aspen species can readily hybridize and their artificial hybrids show heterosis for many growth and wood characteristics (Hart, De Araujo, Thomas, & Mansfield, 2013), suggesting that the speciation process has not gone to completion among the three aspen species. In comparison, *P. trichocarpa* belongs to a different section of the genus *Populus* (section *Tacamahaca*), and it is reproductively isolated from all aspen species (Jansson & Douglas, 2007). Facilitated by the availability of a high-quality reference genome of *P. trichocapra* (Tuskan et al., 2006), the four *Populus* species represent a promising model system to investigate how various evolutionary forces have shaped the evolution of the genomic landscape of differentiation across the speciation continuum in forest trees.

**Figure 1.**
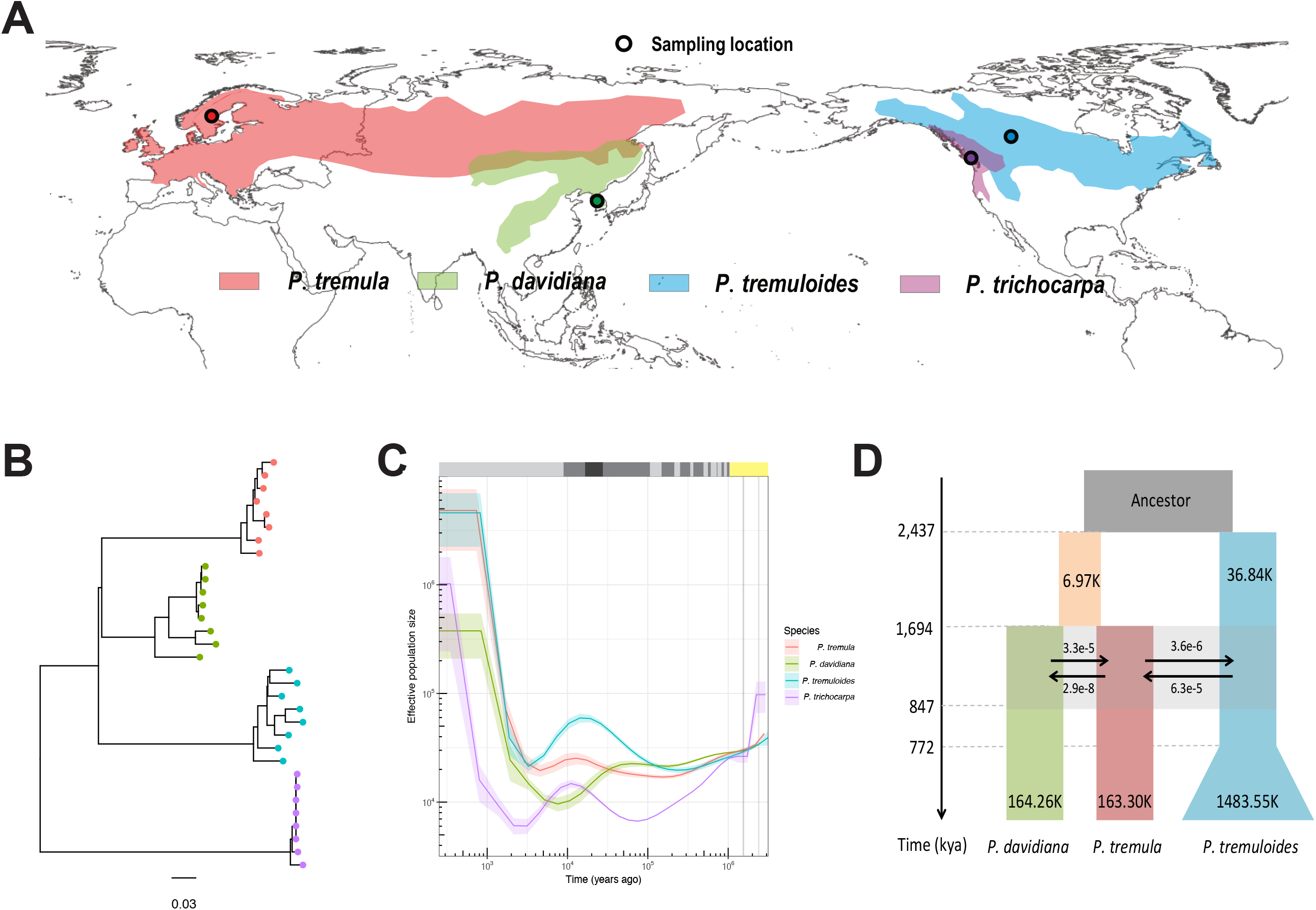
Phylogenetic and population genetic analyses of four *Populus* species. (A) Sampling locations (black circle) of eight individuals from each of the four *Populus* species included in this study. Species ranges for *P. tremula*, *P. davidiana*, *P. tremuloides* and *P. trichocarpa* are indicated by red, green, blue and purple shading, respectively. (B) Maximum-likelihood phylogenetic tree reconstructed based on complete chloroplast sequences. Color scheme for the four species is the same in A-C. (C) Historical effective population size of the four *Populus* species inferred using MSMC v2 based on sets of eight haplotypes, with solid lines representing medians and shading representing ± standard deviation calculated across pairs of haplotypes. Yellow bar indicates Early Pleistocene cooling; Glacial and interglacial periods of the Late and Middle Pleistocene are indicated by dark and light grey bars, respectively; black bar indicates the period of Last Glacial Maximum (LGM). (D) Schematic of demographic scenarios of the three aspen species modeled using fastsimcoal2. The ancestral population is shown in light and dark grey respectively for different ancestral lineages. *P. tremula* is in red, *P. davidiana* is in green, and *P. tremuloides* is in blue. The arrows indicate the per generation migration rate (*m*) between species. Estimated divergence time, effective population size, and gene flow are detailed in Supplementary Table S7.

We use whole-genome re-sequencing in the four *Populus* species to (i) determine their speciation history and characterize whether there is historical gene flow between the now-allopatric species; (ii) examine the fine-scale genomic landscapes of diversity and divergence across species at different stages of divergence; (iii) quantify the extent of genome-wide genealogical discordance and ancient hybridization among the three closely related aspen species; (iv) identify the signatures of positive selection and long-term balancing selection along the genome, and uncover how they impact levels of variation during speciation. Overall, our main aim is to disentangle and understand how the multitude of evolutionary processes have shaped the genomic architecture during speciation.

## 2. Materials and Methods

### 2.1 Sample collection, whole-genome resequencing and genotype calling

We used whole genome resequencing data from eight individuals each of *Populus tremula*, *P. tremuloides* and *P. trichocarpa*, as described in Wang et al. (2016a), and additional eight individuals of *P. davidiana* that are first reported in this study (Table S1). The sampling was from a single geographic region for each species (Figure 1A). Briefly, sequencing of all samples was carried out on the Illumina HiSeq 2000 platform. Prior to read mapping, we used Trimmomatic (Lohse et al., 2012) to remove adapter sequences and to trim low quality bases from the start or the end of reads (base quality ≤ 20). If the processed reads were shorter than 36 bases after trimming, the entire reads were discarded. After quality control, we mapped the remaining reads from each individual to the *P. trichocarpa* reference genome (v3.0) (Tuskan et al., 2006) using BWA-MEM algorithm with default parameters, as implemented in bwa-0.7.10 (Li, 2013).

To minimize the influence of mapping bias, several further filtering steps were employed before genotype calling. First, we used RealignerTargetCreator and IndelRealigner in GATK v3.8.0 (DePristo et al., 2011) to correct for the mis-alignment of bases in regions around insertions and/or deletions (indels). Second, to account for the artifacts due to PCR duplication introduced during library construction, we used the MarkDuplicates method from Picard packages (http://broadinstitute.github.is/picard/) to only retain the read or read-pair with the highest summed base quality among those with identical external coordinates and same insert lengths. Additionally, we further discarded site types that likely cause mapping bias based on three criteria: (1) those with extreme read coverage (less then 4× or higher than twice of the mean coverage); (2) covered by more than two reads of mapping score equaling zero per individual; (3) overlapping known repetitive elements as identified by RepeatMasker (Tarailo - Graovac & Chen, 2009). Finally, sites that passed all these filtering criteria were used in downstream analyses. This left a total of 168,950,389 sites for further analysis (42.8% of collinear genomic sequences of the *P. trichocarpa* genome assembly).

After filtering, we implemented two complementary approaches for genotype calling. First, to account for the bias inherent in genotype calling approach from next generation sequencing (NGS) data (Nielsen, Korneliussen, Albrechtsen, Li, & Wang, 2012), the population genetic estimates that relied on site frequency spectrum (SFS) were calculated using ANGSD v0.917 (Korneliussen, Albrechtsen, & Nielsen, 2014). Second, for the analyses that require accurate single nucleotide polymorphism (SNP) calls, genotype calling in each individual was performed using HaplotypeCaller of the GATK v3.8.0, and GenotypeGVCFs was then used to merge multi-sample records from the four species together for re-genotyping and re-annotation of the newly merged VCF (DePristo et al., 2011). To minimize genotype calling bias and to retain high-quality SNPs, we further performed several filtering steps: (1) SNPs that overlapped with sites not passing all previous filtering criteria were removed; (2) only bi-allelic SNPs with a distance of at least 5 bp away from any indels were retained; (3) genotypes with read depth (DP) < 5 and/or with genotype quality score (GQ) < 10 were treated as missing, and we then removed all SNPs with a genotype missing rate > 10%. After all these steps of filtering, a total of 8,568,990 SNPs were retained across the four *Populus* species. For the analyses that required imputed and phased dataset, BEAGLE v4.1 (Browning & Browning, 2009) was used to infer haplotypes of individuals within each species.

### 2.2 Phylogenetic relationships and population structure analysis

#### Chloroplast phylogeny

To infer the phylogenetic relationship of the four *Populus* species based on chloroplast data, we first mapped the filtered reads from our resequencing data against the *P. trichocarpa* chloroplast genome using bwa-aln 0.7.10 (Li & Durbin, 2009). Then, UnifiedGenotyper in GATK v3.8.0 was used to call SNPs at all sites (--output_mode EMIT_ALL_SITES). Since chloroplasts are haploid and SNPs are thus expected to be homozygous, the haploid option (-ploidy 1) in UnifiedGenotyper was used. After treating sites with GQ < 30 as missing data, only bi-allelic SNPs with quality by depth (QD) ≥ 10 and with a missing rate ≤ 20% were retained. Finally, a consensus tree was constructed based on 1,292 chloroplast SNPs using maximum likelihood method implemented in SNPhylo (Lee, Guo, Wang, Kim, & Paterson, 2014).

#### Principle component analysis (PCA)

To account for the uncertainty in genotype calls, PCA was performed using ANGSD v0.917 and ngsTools v1.0.1 (Fumagalli, Vieira, Linderoth, & Nielsen, 2014). We first used the SAMTools model (Li et al., 2009) in ANGSD to estimate genotype likelihoods from BAM files using only reads with a minimal base quality score of 20 and a minimal mapping quality score of 30 across all individuals. ngsTools was then used to compute the expected covariance matrix across pairs of individuals for the four species based on the genotype posterior probabilities across all filtered sites. Eigenvectors and eigenvalues were generated with the R function eigen from the covariance matrix, and the significance level was determined using the Tracy-Widom test as implemented in EIGENSOFT version 6.1.4 (Patterson, Price, & Reich, 2006).

#### Identity-by-descent (IBD) blocks analysis

To determine the extent to which individuals across the four species shared DNA segments, the identity-by-descent block analysis was performed for the four species using BEAGLE v4.1 (Browning & Browning, 2013) with the following parameters: window=100,000; overlap=10,000; ibdtrim=100; ibdlod=5.

### 2.3 Demographic history reconstruction

#### MSMC

We used Multiple Sequentially Markovian Coalescent approach (MSMC v2) (Schiffels & Durbin, 2014) to infer patterns of historical patterns of effective population sizes changes through time for all four *Populus* species. Only sites passing all above filtering criteria were included in analyses. Because different number of individuals and haplotypes provides different resolution for recent and distant population histories, we applied MSMC to phased whole-genome sequences from one (two haplotypes, which can infer more distant size changes), two (four haplotypes, which infer size changes at intermediate time scales) and four (eight haplotypes, which infer the most recent size changes) individuals for each species, respectively. We did not include more haplotypes due to the computational cost of using larger haplotype sets. In total, we have 8, 28 and 70 different individual configurations for two-, four-, and eight-haplotype analyses in each species. We ran MSMC on all individual configurations and estimated medians and standard deviations of effective population sizes changes across time. To convert the coalescent scaled time to absolute time in years, we used a mutation rate of 2×10^-9^ per site per year (Koch, Haubold, & Mitchell-Olds, 2000) and a generation time of 15 years.

#### Fastsimcoal2

Given the long divergence time and the low number of polymorphic sites shared between aspens and *P. trichocarpa* (Wang, Street, Scofield, & Ingvarsson, 2016a), we used a coalescent simulation-based method implemented in *fastsimcoal2.6* (Excoffier, Dupanloup, Huerta-Sánchez, Sousa, & Foll, 2013) to only infer the demographic and speciation histories only for the three aspen species. For all possible pairs of the three species, the two-dimensional joint SFS (2D-SFS) was constructed from the posterior probabilities of sample allele frequencies using ANGSD v0.917 (Figure S1). A total of twenty-nine models were evaluated and all models began with the split of an ancestral population into the Eurasia and the North America lineage (*P. tremuloides*) followed by the split of the Eurasian lineage into *P. tremula* and *P. davidiana*. The models differed in terms of (1) whether post-divergence gene flow was present or not, (2) time, level and pattern of gene flow between the three aspen species, and (3) the occurrence and pattern of population expansion in *P. tremuloides* given a genome-wide excess of rare frequency alleles that we observed in this species in our previous study (Wang et al., 2016a) and also in this study (Figure S2). Alternative demographic models were fitted to the joint SFS data. The global maximum likelihood estimates for all demographic parameters under each model were obtained from 50 independent runs, with 100,000 coalescent simulations per likelihood estimates (-n 100000, -N 100000) and 40 cycles of the likelihood maximization algorithm. The models were compared based on the maximum value of likelihood over the 50 independent runs using the Akaike’s weight calculated following Excoffier et al. (2013). The model with the maximum Akaike’s weight value was chosen as the optimal one. Confidence intervals were generated by performing parametric bootstrapping with 100 bootstrap replicates, and with 50 independent runs in each bootstrap. As for MSMC, we assumed a mutation rate of 2×10^-9^ per site per year and a generation time of 15 years (Koch et al., 2000) when converting estimates to units of years and individuals.

### 2.4 Intra- and inter- species summary statistics

#### Intra-species genomic diversity

For each species, based on the SAMTools genotype likelihood model (Li et al., 2009), we used ANGSD v0.917 (Korneliussen et al., 2014) to estimate allele frequency likelihoods, obtain a maximum likelihood estimate of the folded site frequency spectrum and used this to calculate nucleotide diversity (π) in non-overlapping sliding windows of 10 Kbp and 100 Kbp across the entire genome. Only sites with a minimum mapping quality of 30 and minimum base quality of 20 were used in the estimation. Windows were discarded if there were less than 10 % sites left after all of the filtering steps described above. Since our previous study showed that linkage disequilibrium (LD) decays within 10 Kbp in different species of *Populus* (Wang et al., 2016a), in the following we focused more on estimates derived from 10 Kbp windows.

#### Inter-species genomic divergence

For each species pair, we estimated two divergence metrics across the 10 Kbp and 100 Kbp non-overlapping windows: genetic differentiation (*F*_ST_) and sequence divergence (d_xy_). Without relying on SNP or genotype calling (Fumagalli et al., 2013), we first used ANGSD to calculate posterior probabilities of sample allele frequency for each species. Then, the program ngsFST from the ngsTools package was used to estimate *F*_ST_ between species using a method-of-moments estimator, and the program ngsStat was used to calculate d_xy_ between species at each site. Finally, we averaged these divergence values across all sites within each window.

#### Population-scaled recombination rate

For each species we used LDhelmet v1.9 (Chan, Jenkins, & Song, 2012), a coalescent-based, reversible-jump Markov chain Monte Carlo (rjMCM) simulation method, to estimate the population-scaled recombination rate, ρ. First, VCFtools (Danecek et al., 2011) and custom shell script were used to tailor the phased genotype data of each chromosome to the necessary input sequence file (fasta format). Then, we used ‘find_confs’ in LDhelmet to concatenate all the input sequences files and generate a haplotype configuration file per species. Thereafter, ‘table_gen’ was used to compute the likelihood lookup table for each species, where we assume the approximate genome-wide neutral diversity (θ) of 0.01 for the three aspen species and of 0.005 for *P. trichocarpa* (Wang et al., 2016a), and the grid of ρ values was specified as –r 0.0 0.1 10.0 1.0 100.0 for all species. In addition, the optional ‘pade’ component of LDhelmet was included in the analysis, which computes the Padé coefficients (-x 11) from the haplotype configuration file. Finally, we ran LDhelmet with window size of 50 SNPs and block penalty of 50 for a total of 1,000,000 iterations, discarding the first 100,000 as burn-in. We then calculated weighted average of the estimated ρ in 10 Kbp and 100 Kbp windows, respectively. Windows with less than 50 SNPs (for 10 Kbp windows) and 200 SNPs (for 100 Kbp windows) left from previous filtering steps were discarded.

### 2.5 Window-based phylogenomic analysis

#### Topology weighting

As expected for a clade with rapid radiation, genealogies may vary widely across different genomic regions (Lamichhaney et al., 2015). Given that *P. trichocarpa* is distantly related from the other three aspen species (Hamzeh & Dayanandan, 2004), we used Twisst, a topology weighting method by iterative sampling of subtrees (Martin & Van Belleghem, 2017), to assess and quantify the phylogenetic discordance among the three aspen species along the genome. The genealogical relationships of these species can be defined by three possible topologies: [(*P. tremula*, *P. davidiana*), *P. tremuloides*], [(*P. tremula*, *P. tremuloides*), *P. davidiana*], [(*P. davidiana*, *P. tremuloides*), *P. tremula*]. Using *P. trichocarpa* as the outgroup species, local phylogenetic subtrees was inferred in RAxML v8.2.4 (Stamatakis, 2014) with the GTRCATI model over non-overlapping 10 Kbp and 100 Kbp windows. Topology weightings for each window were then computed through determining the number of unique subtrees that match each of the three possible topologies by iteratively sampling a single haplotype from each species (Martin & Van Belleghem, 2017). Windows were discarded in topology weighting estimation if there were < 50 SNPs and < 200 SNPs left from previous filtering steps for 10 Kbp and 100 Kbp windows, respectively.

#### Inference of incomplete lineage sorting

Because the speciation events that resulted in aspen species were close in time (see Results), we expect the lineage sorting process relating these species to be incomplete. Given the three aspen species and the outgroup poplar species (*P. trichocarpa*) with the relationship as (((*P. tremula*, *P. davidiana*), *P. tremuloides*), *P. trichocarpa*), we labeled alleles in *P. tremula*, *P. davidiana* and *P. tremuloies* as A (ancestral allele) if they match the reference allele of *P. trichocarpa* genome, and B (derived allele) otherwise. We then considered segregating sites with (((*P. tremula*, *P.davidiana*), *P. tremuloides*), *P. trichocarpa*) patterns as AABAs, ABAAs, BAAAs, ABBAs, BABAs and BBAAs. The two SNP patterns ABBAs and BABAs can result from incomplete lineage sorting if we assume no gene flow occurred among species (Durand, Patterson, Reich, & Slatkin, 2011; Green et al., 2010). We calculated the level of incomplete lineage sorting (ILS) at site *i* in the genome as:

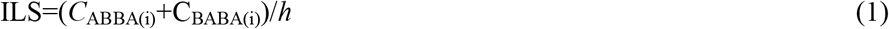

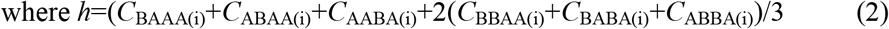

Because population samples were used for all species, at each site we used the frequency of the derived allele in each species to effectively weight each segregating site according to its fit to the six segregation patterns for the three aspen species (Durand et al., 2011), with

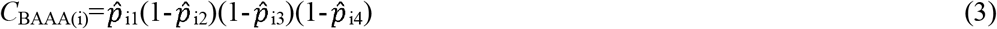

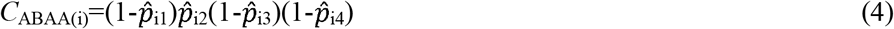

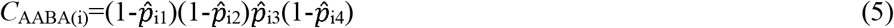

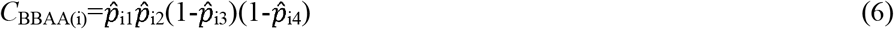

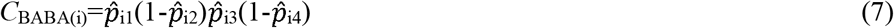

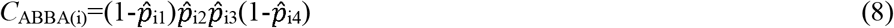

where 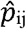 is the frequency of the derived allele at site i in species j.

The calculation of ILS presents the counts of incomplete lineage sorting pattern (*C*_ABBA(i)_ and C_BABA(i)_) normalized by the total count of segregating sites (*h*), which is a proxy of the species tree topology height (Scally et al., 2012). We then summarized the proportion of ILS over non-overlapping 10 Kbp and 100 Kbp windows or in bins with varying distances from the nearest exon.

### 2.6 Analyses of introgression

We first tested for introgression between the three aspen species using the *D*-statistic, also known as the ABBA-BABA tests. These tests evaluate the imbalance frequency of site patterns (Durand et al., 2011; Green et al., 2010). Using *P. trichocarpa* as the outgroup, we expect equal counts of the two site patterns (ABBA and BABA) when incomplete lineage sorting causes the site pattern discordance. If discordance is caused by introgression, one of the site patterns is expected to be more prevalent than the other. We applied two approaches to perform the *D*-statistic test. First, we used –doAbbababa2 implemented in ANGSD v0.917 (Korneliussen et al., 2014) to directly count ABBA and BABA sites and calculate the *D* statistic without calling genotypes in non-overlapping 10 Kbp and 100 Kbp windows for the whole genome. Then, jackknife bootstrapping was conducted to estimate significance at the chromosome and whole-genome level. Second, based on allele frequencies at each SNP called by GATK, we calculated the *D* statistic in 10 Kbp and 100 Kbp non-overlapping windows across the genome with the python script *ABBABABAwindows.py* (https://github.com/simonhmartin/genomics_general) (Martin, Davey, & Jiggins, 2014). Following the detection of introgression among individuals at the genome level, we used a modified *f*-statistic (*f*_d_) (Martin et al., 2014) to estimate the proportion of introgressed sites at the population level using *ABBABABAwindows.py* (https://github.com/simonhmartin/genomics_general) on non-overlapping 10 Kbp and 100 Kbp windows across the genome.

### 2.7 Genome-wide scan for regions under positive and balancing selection in aspens

To specifically test for the impact of positive and balancing selection on the genomic landscape during speciation of aspens, we first used a composite likelihood ratio (CLR) statistic implemented in SweepFinder2 (DeGiorgio, Huber, Hubisz, Hellmann, & Nielsen, 2016) to detect regions subject to recent positive selection or selective sweeps in each of the three aspen species. The ancestral allelic state was defined by assuming that the alleles that were the same as those found in the *P. trichocarpa* reference genome was the ancestral alleles. By contrasting the likelihood of the null hypothesis, based on the unfolded site frequency spectrum (SFS) calculated across the genome using –f option, with the likelihood of a model where the SFS has been altered by a recent positive selection event, the CLR statistics was calculated in non-overlapping windows of 10 Kbp. Within each species, windows with CLR values higher than the 99^th^ percentile of its distribution were identified as candidate region under selection.

Moreover, we identified regions under long-term balancing selection by estimating the summary statistics, β (beta score), which detects the clusters of variants with an excess number of intermediate frequency polymorphisms (Siewert & Voight, 2017). Given that the signals of long-term balancing selection usually is localized to a narrow genomic region (Gao, Przeworski, & Sella, 2015), we used 1 Kbp windows to calculate β values for each core SNPs in the three aspen species. We used the unfolded version of β, with the ancestral and derived allelic states inferred based on comparisons with the outgroup species *P. trichocarpa*. To prevent false positives, we filtered out SNPs with a folded frequency lower than 20%, and defined the SNPs with extreme β scores in the top 1% as significant. Furthermore, only SNPs that are significant in all three species were considered as putative targets of long-term balancing selection. Finally, we binned significant SNPs into 10 Kbp windows for downstream comparisons.

Lastly, to assess the effects of positive and balancing selection on the genomic architecture of speciation, we compared outlier windows that were identified as being under positive or balancing selection with the remaining genomic regions using a variety of population genetic statistics, including genetic diversity, divergence, lineage sorting and introgression within and between the three closely related aspen species. Differences between outlier windows and the genome-wide averages for all these statistics were tested using Wilcoxon ranked-sum tests. To further examine whether any functional classes of genes were over-represented in these candidate regions, we performed gene ontology (GO) analyses using the R package topGO 2.36.0 (Alexa & Rahnenführer 2009). Fisher’s exact test was used to calculate the statistical significance of enrichment, and GO terms with *P*-value lower than 0.01 were considered to be significantly enriched.

## 3. Results

### 3.1 Phylogenetic relationships, population structure and demographic history

The genome alignment resulted in an average depth of 24.6× across all individuals after quality control (Table S1). The PCA results revealed a clear distinction among the four *Populus* species (Figure S4). Based on the Tracy-Widom test, only the first three components were significant (Table S2). The first principal component (PC1; variance explained=28.79%) separated *P. trichocarpa* from the three aspen species, while the second component (PC2; variance explained=7.52%) separated *P. tremuloides* from *P. tremula* and *P. davidiana*. Finally, the third component (PC3, variance explained=5.33%) separated *P. tremula* and *P. davidiana*. The clustering and genetic relationships of the four species were also confirmed by the phylogenetic tree constructed based on the entire chloroplast genomes (Figure 1B). Moreover, we measured the number and length of shared IBD haplotypes within and between species (Figure S5, S6; Table S3, S4). Compared to between-species comparisons, we found much more extensively shared IBD haplotypes for within-species comparisons (Figure S5), although haplotypes shared within *P. tremuloides* were shorter than the other three species (Figure S6A; Table S4). This is likely owing to the higher recombination rate and more rapid decay of linkage disequilibrium (LD) in *P. tremuloides* than other species (Wang et al., 2016a). For the between-species comparisons, we did no observe any haplotype sharing between the three aspen species and *P. trichocarpa*, confirming the distant relationship between aspens and poplars in the genus *Populus* (Figure S5, Table S3). Within aspens, *P. tremula* and *P. davidiana* shared more and longer haplotypes than either of them shared with *P. tremuloides* (Figure S5, S6B; Table S3, S4), which also supports a closer relationship between these two species, as identified in the chloroplast phylogeny.

To investigate the demographic and speciation histories of the four *Populus* species, we first used MSMC to examine historical fluctuations in the effective population size (*N*_e_) for each species. The results showed that all species experienced a period of population decline during the early Pleistocene cooling (2.5-0.9 million years ago, Mya) (Figure 1C). Compared with the three aspen species, *P. trichocarpa* experienced a more dramatic population decline during this period (Figure 1C, Figure S7), which likely explain the much lower genetic diversity observed in this species relative to others (Table S5). The two North American species, *P. tremuloides* and *P. trichocarpa*, experienced a population expansion from the start of the last ice age (110 thousand years ago, Kya) until the last glacial maximum (LGM, 23-18 Kya) whereas the European species *P. tremula* remained relatively stable. On the other hand, the eastern Asian species, *P. davidiana*, showed pattern of population decline during the entire period (Figure 1C, Figure S7). Our results therefore revealed that before the LGM, forest trees distributed in different continents experienced asynchronous demographic responses to Pleistocene climate changes (Bai et al., 2018). During and following the LGM, all four species experienced a population decline followed by a subsequent rapid population expansion (Figure 1C).

Because of the distant phylogenetic relationship and low levels of shared polymorphism between *P. trichocarpa* and the three aspen species (Wang et al., 2016a), we therefore explicitly focused on inferring the demographic parameters of the speciation history for the three aspens. After evaluating a total of twenty-nine models (Figure S8), the best-supported model (Figure S8; Table S6) suggests that the Eurasian lineage (the common ancestor of *P. tremula* and *P. davidiana*) split from the North American lineage (*P. tremuloides*) at ∼ 2.4 Mya (bootstrap range [BP]: 2.1-3.2 Mya), which is in accordance with our earlier estimates on the divergence time between *P. tremula* and *P. tremuloides* (Wang, Street, Scofield, & Ingvarsson, 2016b). The European lineage (*P. tremula*) and the East-Asian lineage (*P. davidiana*) diverged ∼1.7 Mya (BP: 1.5-2.1 Mya) (Figure 1D, Table S7). Our results detected low levels of ancient gene flow between *P. tremula* and *P. davidiana*, and between *P. tremula* and *P. tremuloides* following speciation until around 847 Kya (BP: 539Kya-1.0 Mya) (Figure 1D). After this period the species have remained isolated which is also reflected by their current disjunct geographic distributions (Figure 1A). Compared to the Eurasian lineage of aspens, *P. tremuloides* has experienced a notable population expansion in the recent past (∼772 Kya, BP: 440-887 Kya), which is consistent with its genome-wide excess of rare alleles (Figure S2).

### 3.2 General patterns of genome-wide diversity and differentiation

We further characterized genome-wide patterns of nucleotide diversity (π), population recombination rate (ρ) and divergence (*F*_ST_ and d_xy_) for the four *Populus* species (Figure 2A; Table S5, Table S8-S10). At the species level, π varied markedly between species, ranging from 0.0063 in *P. trichocarpa* to 0.0148 in *P. tremuloides*, but the average genomic diversity was very similar across the three aspen species (Table S5). In contrast to the patterns observed for π, the population-scaled recombination rate, ρ, was much higher in *P. tremuloides* (0.0273 bp^-1^) than in the other three species (0.0096 bp^-1^-0.0139 bp^-1^) (Table S8). Variation in genetic divergence (*F*_ST_ and d_xy_) among the six species pairs reveals the continuous nature of differentiation along the speciation continuum, with *P. tremula* and *P. davidiana* showing the lowest levels of divergence and with the highest divergence observed between aspens and *P. trichocarpa* (Figure 2A, Table S9, S10).

**Figure 2.**
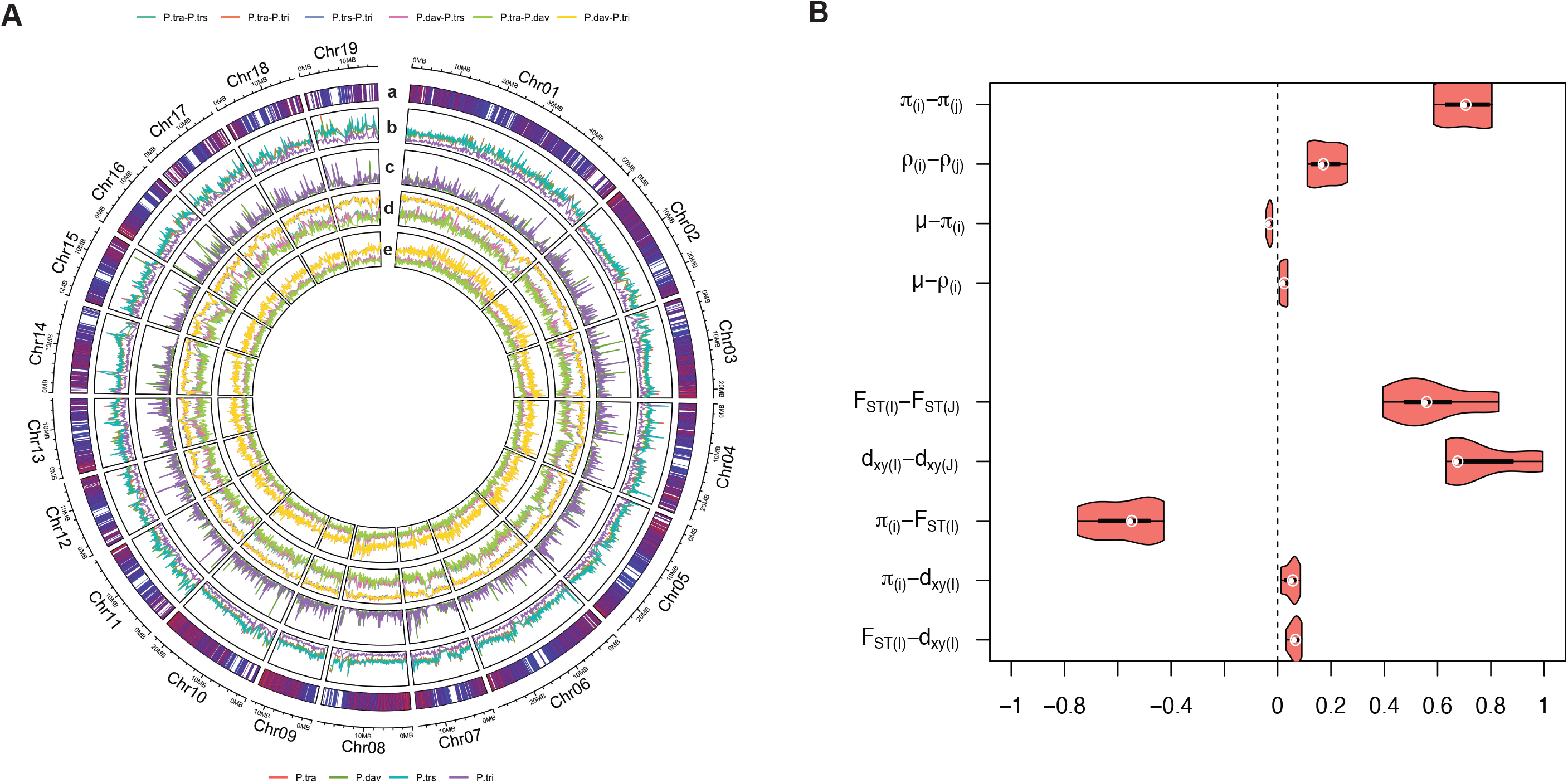
Genome-wide landscape of genetic diversity and divergence within and between species. (A) Chromosomal landscape of (a) the density of coding sequences (CD); (b) nucleotide diversity (π); (c) recombination rate (*ρ*); (d) the relative measure of genetic divergence (*F*_ST_) and (e) the absolute measure of genetic divergence (d_xy_). (B) Distribution of correlation coefficients (Spearman’s ρ) shown as violin plots for population summary statistics characterizing genomic features (neutral mutation rate *μ*) and variation within (π, *ρ*) and between species (*F*_ST_, d_xy_) calculated at 100 Kbp windows. Subscripts ‘i, j’ symbolize all possible combinations of correlations between two species i=1…(n-1) and j=(i+1)…n for within-species measures; Capital letters ‘I, J’ symbolize inter-species statistics. Correlations exclude pseudo-replicated species comparisons. Detailed information can be found in Supplementary Table S11.

At the genome level, patterns of genetic diversity and divergence show high levels of parallelism in all pairwise comparisons. The genome-wide profiles of π (average Spearman’s *ρ* =0.71) and ρ (average Spearman’s *ρ* =0.18) were positively correlated in all possible species pairs (Figure 2B, Table S11). We found little evidence for an association between either π or ρ and the local mutation rate (μ, approximated by the four-fold synonymous substitution rate) (Figure 2B, Table S11). Hence, the broad-scale variation in genetic diversity is conserved across the diverging lineages, which likely arise from a common genomic architecture where linked selection has played a major role in shaping local genomic diversity (Burri, 2017). This is further highlighted by the conserved landscape of recombination rate variation across the genomes of the species and the strong degree of genome synteny that we observed between the genomes of aspens and poplars (Lin et al., 2018). Second, we found that the differentiation landscapes were highly correlated among all species pairs both for the relative (*F*_ST_) and the absolute (d_xy_) measures of genetic differentiation (Figure 2B, Figure S9, Table S11). The highly similar landscape of differentiation among different species pairs could imply phylogenetically conserved genomic features, e.g. conserved landscapes of functional densities and recombination (Burri, 2017; Vijay et al., 2017). Moreover, significantly negative correlations between *F*_ST_ and π were found in all pairwise comparisons (Figure 2B, Figure S9, Table S11), which is in line with the observation that *F*_ST_ is sensitive to intra-specific nucleotide diversity (Charlesworth, 1998). In contrast, only weak correlations were observed between d_xy_ and π. Because d_xy_ largely reflects diversity in a common ancestor (Cruickshank & Hahn, 2014), a weak correlation between d_xy_ and π implies that ancestral diversity might have little impact on extant diversity in the different *Populus* species. In addition to extant diversity, d_xy_ was only weakly correlated with *F*_ST_ across all comparisons (Figure 3B, Figure S9, Table S11), which further implies that ancestral polymorphisms have had limited contribution to the genomic divergence of extant species (Cruickshank & Hahn, 2014).

**Figure 3.**
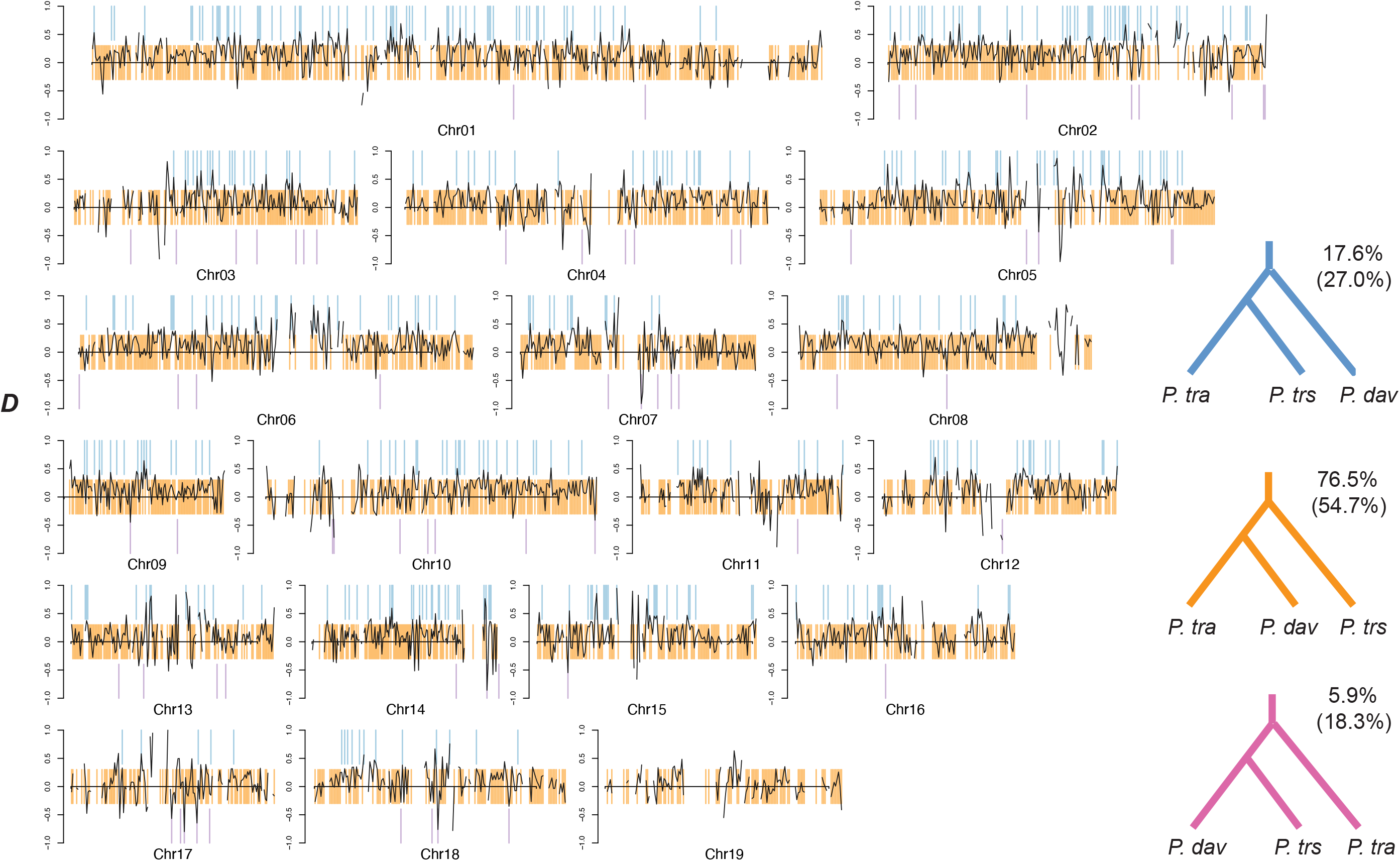
Heterogeneous distribution of phylogenies in three aspen species. Chromoplots for 19 chromosomes show the distribution of three possible rooted phylogenetic relationships inferred from 100 Kbp genomic regions for *P. tremula* (*P. tra*), *P. davidiana* (*P. dav*) and *P. tremuloides* (*P. trs*), with *P. trichocarpa* as outgroup species. The colored vertical bars represent the windows with complete monophyly of the three alternative topologies as shown in the lower right, where the proportion of the three topologies in 100 Kbp and 10 Kbp (in parenthesis) across the genome are also shown. Across all chromosomes, the *D* statistic generally tends toward positive values, indicating ancient introgression between *P. tra* and *P. trs* may have been occurring across the genome.

### 3.3 Topology weighting reveals phylogenetic discordance and ancient introgression between P. tremula and P. tremuloides

Even if the analyses of current population structure and genomic divergence support a clear species relationship for the four *Populus* species, (((*P. tremula*, *P. davidiana*), *P. tremuloides*), *P. trichocarpa*), we used a topology weighting approach to explore to what degree the ‘species tree’ was congruent across the entire genome. Using *P. trichocarpa* as an outgroup, our results reveal widespread incongruence in local genealogies in either 10 Kbp or 100 Kbp non-overlapping windows across the genome (Figure 3, Figure S10). The most prevalent topology, ((*P. tremula*, *P. davidiana*), *P. tremuloides*), which reflects the likely ‘species topology’, has an average weighting of 54.7% and 76.5% across the genome in 10 Kbp and 100 Kbp windows, respectively. Of the other two topologies, the ((*P. tremula*, *P. tremuloides*), *P. davidiana*) topology was much more common (27.0% and 17.6% for 10 Kbp and 100 Kbp windows) compared to ((*P. davidiana*, *P. tremuloides*), *P. tremula*) (18.3% and 5.9% for 10 Kbp and 100 Kbp windows) (Figure 3, Table S12). In general, we observed that larger windows (100 Kbp) produced higher rates of monophyly (windows with a weighting of 1) and a greater fraction of resolved trees compared to the smaller windows (10 Kbp) (Figure S10, Table S13).

Interestingly, in contrast to all other chromosomes where all three topologies were observed, chromosome 19, which is known to harbor the sex determination region in *Populus* (Yin et al., 2008), showed only a single monophyletic grouping of the ‘species topology’ (Figure 3). Such a pattern is consistent with the expectation that lineage sorting is faster on sex chromosomes compared to autosomes because of its smaller effective population size (Meisel & Connallon, 2013; Vicoso & Charlesworth, 2006). Overall, both incomplete lineage sorting (ILS) and introgression can result in discordance between the local topology and the species tree for recently diverged species. Given that ILS is expected to generate equal frequencies of the alternative topologies (Durand et al., 2011; Mailund et al., 2014), the more frequent topology of ((*P. tremula*, *P. tremuloides*), *P. davidiana*) is likely explained by the occurrence of introgression between *P. tremula* and *P. tremuloides*. We therefore compared the distribution of the branch lengths separating each pair of aspen species among all topology types. Compared to the expectation that species with recent introgression tend to be separated by short branches (Fontaine et al., 2015; Martin & Van Belleghem, 2017), the branch distances between *P. tremula* and *P. tremuloides* were not obviously different from other species-pairs across topology comparisons (Figure S11). This pattern is most likely caused by ancient hybridization between these two species where genetic drift has eradicated most signatures of gene flow after an ancient introgression event (Schumer, Cui, Powell, Rosenthal, & Andolfatto, 2016).

To further investigate patterns of ancient introgression between *P. tremula* and *P. tremuloides*, we calculated two statistics associated with the ABBA-BABA test across the genome. The *D*-statistic is used to test for ancient gene flow by comparing the imbalance of ABBAs and BABAs, and the *f*_d_-statistic is used to estimate the fraction of the genome shared through ancient introgression. For the *D*-statistics, we also implemented two different approaches, which differed in whether the called genotypes was relied or not. We find that the estimates of the two approaches are highly correlated with each other (Figure S12), suggesting that this statistic is robust to identify introgression regardless of which type of data is used. Genome-wide estimates of the *D*-statistic and *f*-statistic showed a general pattern of positive values over 10 Kbp and 100 Kbp non-overlapping windows (Table S14), confirming that *P. tremuloides* has a closer genetic relationship with *P. tremula* than with *P. davidiana*. Thus, the significant asymmetry in genetic relationship together with the excess of shared sequence polymorphism between *P. tremula* and *P. tremuloides* (Figure S13) all provide evidence for historical gene flow between the currently allopatric Eurasian and North American aspen species.

### 3.4 Long-term effects of selection in shaping patterns of diversity, divergence, incomplete lineage sorting (ILS) and levels of introgression in Populus species

To evaluate the impact of natural selection on genetic diversity, divergence, ILS and gene flow in the context of speciation, we examined the correlations between these genetic parameters and factors affecting the extent and efficiency of selection. First, regions with a high density of potential targets for selection are expected to experience stronger linked selection simply because selection occurs more often in such regions (Al-Shahrour et al., 2010; Flowers et al., 2011). We therefore examined the relationship between intraspecific diversity, species divergence and the density of functional elements, defined as the proportion of protein-coding sites within a 10 Kbp or 100 Kbp window (coding density). We hypothesized that if selection contributes to the reduction of diversity at linked neutral sites, its effect is expected to be more pronounced in regions with greater content of functional elements (Ravinet et al., 2017). Consistent with this prediction, we observed a significantly negative relationship between π and functional content (Figure 4A). This correlation was robust to the presence of confounding factors such as GC content, recombination rate and the choice of window size (Table S15).

**Figure 4.**
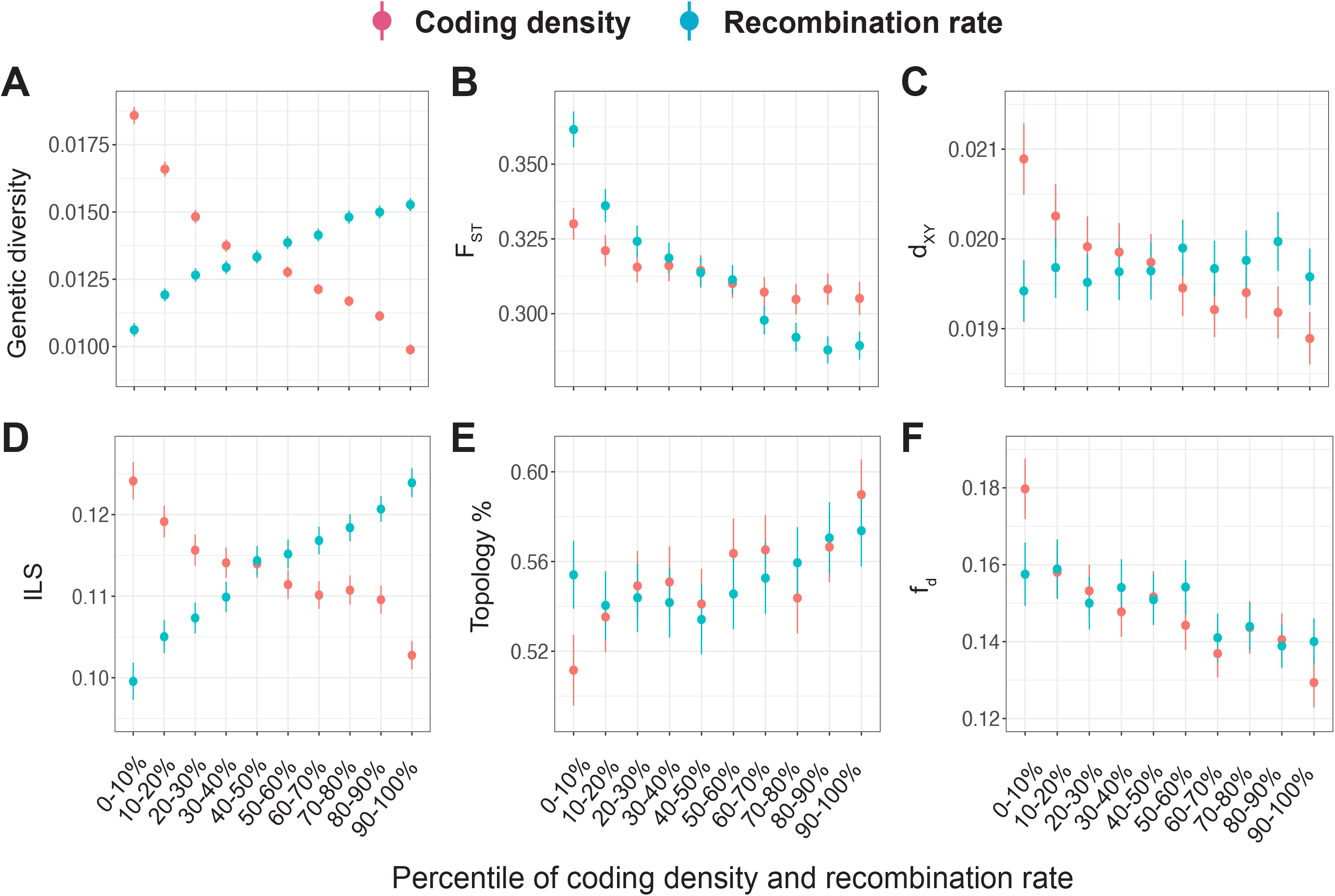
Widespread impact of linked selection. Relationship between recombination rate (blue), coding density (red) and (A) genetic diversity, (B) *F*_ST_, (C) d_xy_, (D) incomplete lineage sorting (ILS), (E) weighting of the ‘species’ tree ([*P. tra*, *P. dav*], *P. trs*)) and (F) the estimated admixture proportion (*f*_d_) between *P. tra* and *P. trs*. Quantile binning is for visualization. The points and error bars indicate the means and 1.96×standard errors. Statistical tests were performed on the unbinned data and detailed correlation coefficients are shown in supplementary Table S15-Table S22.

Moreover, if natural selection was acting on the ancestral polymorphisms prior to the divergence of the two descendant lineages, it could also have an effect on the genetic divergence between species (Munch et al., 2016; Scally et al., 2012). We therefore examined the relationship between interspecies divergence (both *F*_ST_ and d_xy_) and coding density, and observed negative relationships for both *F*_ST_ and d_xy_ (Figure 4B, C), especially between species with longer divergence times (e.g. aspens and *P. trichocarpa*) (Table S17, Table S19). Indeed, if a region experiences natural selection during divergence, it should show lower π within species and higher *F*_ST_ between species because *F*_ST_ is sensitive to intra-specific genetic variation (Cruickshank & Hahn, 2014). Accordingly, a positive correlation between coding density and *F*_ST_ is predicted. The opposite pattern we observe here indicates that long-term natural selection, most likely due to background selection in functional regions, has continuously contributed to the reduced ancestral polymorphism and genetic divergence in regions with greater functional content (Phung et al., 2016). In fact, because of the accumulation of the large amount of new mutations since speciation, ancestral polymorphism may only account for a small amount of the overall average divergence between distantly related species (Edwards & Beerli, 2000). However, the variance of ancestral polymorphism, largely affected by natural selection in ancestral populations, can on the other hand make a substantial contribution to the variability of genome-wide patterns of divergence between species (McVicker, Gordon, Davis, & Green, 2009; Phung et al., 2016).

To further explore the role of natural selection during the divergence of the three aspen species, we examined the extent of ILS across the genome, which can aid to infer evolutionary process in ancestral populations (Mailund et al., 2014; Pease & Hahn, 2013). The pattern of ILS along the genome offers information about the local differences in the ancestral effective population size of the *aspen* ancestor (Pamilo & Nei, 1988). Both purifying and positive selection in the ancestral population are expected to reduce ancestral population size in regions targeted by selection, resulting in increased rates for lineages to coalesce and leaving less available for ILS (Dutheil, Munch, Nam, Mailund, & Schierup, 2015; Munch et al., 2016; Prüfer et al., 2012; Scally et al., 2012). In agreement with this, we found that the fraction of ILS decreases with increasing coding density (Figure 4D), and this relationship remained even after correcting for the confounding variables (Table S21). Within coding exons, ILS is ∼19 % lower and the suppression of ILS extends several thousand bps away from coding genes (Figure S14). Similarly, the proportion of the topology reflecting the true species tree increases with coding density (Figure 4E; Table S21).

In addition, given that the level of admixture estimated by *f*_d_ between *P. tremula* and *P. tremuloides* show considerable heterogeneity across the genome (Figure S15), we examined whether selection may have played a role in shaping genome-wide patterns of introgression. We estimated the relationship between *f*_d_ and coding density and found a significantly negative correlation (Figure 4F; Table S21), indicating that there is greater selection against introgressed alleles in regions enriched for genes (Harris & Nielsen, 2016). The occurrence of this pattern is not likely an artefact of reduced power, as regions with a high density of functionally important elements are expected to have experienced stronger long-term selection and exhibit lower levels of ILS. Accordingly, our power to detect introgression is expected to be elevated close to these regions (Sankararaman et al., 2014; Sankararaman, Mallick, Patterson, & Reich, 2016). Taken together, it is clear that natural selection has had a strong impact on patterns of phylogenetic discordance across the genome among closely related aspen species. However, it is not yet clear to what extent this heterogeneity might be due to incomplete lineage sorting of ancestral polymorphisms or due to ancient introgression. More explicit experimental designs in future studies are needed to tease apart these different processes and explore how natural selection and hybridization act in combination to shape the genome-wide phylogenetic heterogeneity among recently diverged species.

In addition to the local density of functional elements, recombination rates can also interact with natural selection to influence the genomic distribution of genetic diversity and divergence (Figure 4). High recombination can rapidly decouple linked loci and restrict the effect of selection on linked neutral sites (Begun & Aquadro, 1992; Cutter & Payseur, 2013). We found that π and *F*_ST_ showed positive and negative correlations, respectively, with local recombination rates (Figure 4A-C; Table S16, S18, S20). In contrast to the predicted pattern of speciation with gene flow where reduced d_xy_ is expected in regions of high recombination (Nachman & Payseur, 2012), we did not find any relationship between d_xy_ and recombination rate. These observations are in accordance with the expectation that linked selection is prevalent and has genome-wide effects in shaping patterns of genetic diversity and divergence at linked sites in *Populus* (Nachman & Payseur, 2012; Wang et al., 2016b). On the other hand, we found that the incidence of ILS increases with the recombination rate (Figure 4D), which was still observable even after correcting for the confounding variables of coding density and GC content (Table S22). Given that variation in ILS across the genome approximately reflects variation in ancestral *N*_e_ (Degnan & Salter, 2005; Pamilo & Nei, 1988), the stronger effects of recurrent natural selection in low-recombination regions also reduced *N*_e_ in ancestral populations and hence made ILS less likely to occur (Charlesworth et al., 1993; Martin, Davey, Salazar, & Jiggins, 2019; Pease & Hahn, 2013). We did not find obvious correlation between *f*_d_ and recombination rate (Figure 4F, Table S22), might because barriers to introgression has been sculpted by long-term selection and genetic drift after the ancient gene flow and cannot be predicted by recombination rate estimated from current populations. Overall, all these results suggest that the patterns of diversity, divergence and genealogical relationships among the three closely related aspen species are not randomly distributed along the genome, but instead are strongly structured by the interaction between widespread natural selection and intrinsic genomic features, as well as their influence on retention of signatures of ancient gene flow.

### 3.5 The impact of positive and balancing selection on genomic architecture of speciation

Although widespread background selection is likely to have had a large effect in shaping the heterogeneous genomic landscape of variation within and between species (Burri, 2017; Charlesworth, 2012), we were interested in assessing whether positive selection or long-term balancing selection have also played important roles in driving these processes. To identify the impact of positive selection, we performed a composite-likelihood based (CLR) test to scan the genomes for signals of positive selection in each of the three aspen species (Figure 5A). For each species, we considered the windows with a CLR value in the top 1 percentile as candidate region under positive selection. In total, we detected 538 outlier windows across the three species, and only 13 among them were shared by all species (Figure 5B). Our results suggest that most putative sweeps are likely species-specific and may result from relatively recent positive selection that has occurred independently in various lineages after speciation. Compared to genome-wide averages, outlier windows have significantly lower nucleotide diversity, lower recombination rates, higher *F*_ST_ but similar d_xy_ (Figure 5C). In addition, the outlier windows show significantly higher average weightings of the ‘species topology’ (Topo2) and lower levels of ILS compared to genomic background (Figure 5C,D). The ancestral admixture proportion (*f*_d_) between *P. tremula* and *P. tremuloides* is also significantly reduced in the outlier windows (Figure 5D), suggesting that strong selection in these regions may have contributed to the reproductive barriers isolating closely related species (Martin et al., 2019).

**Figure 5.**
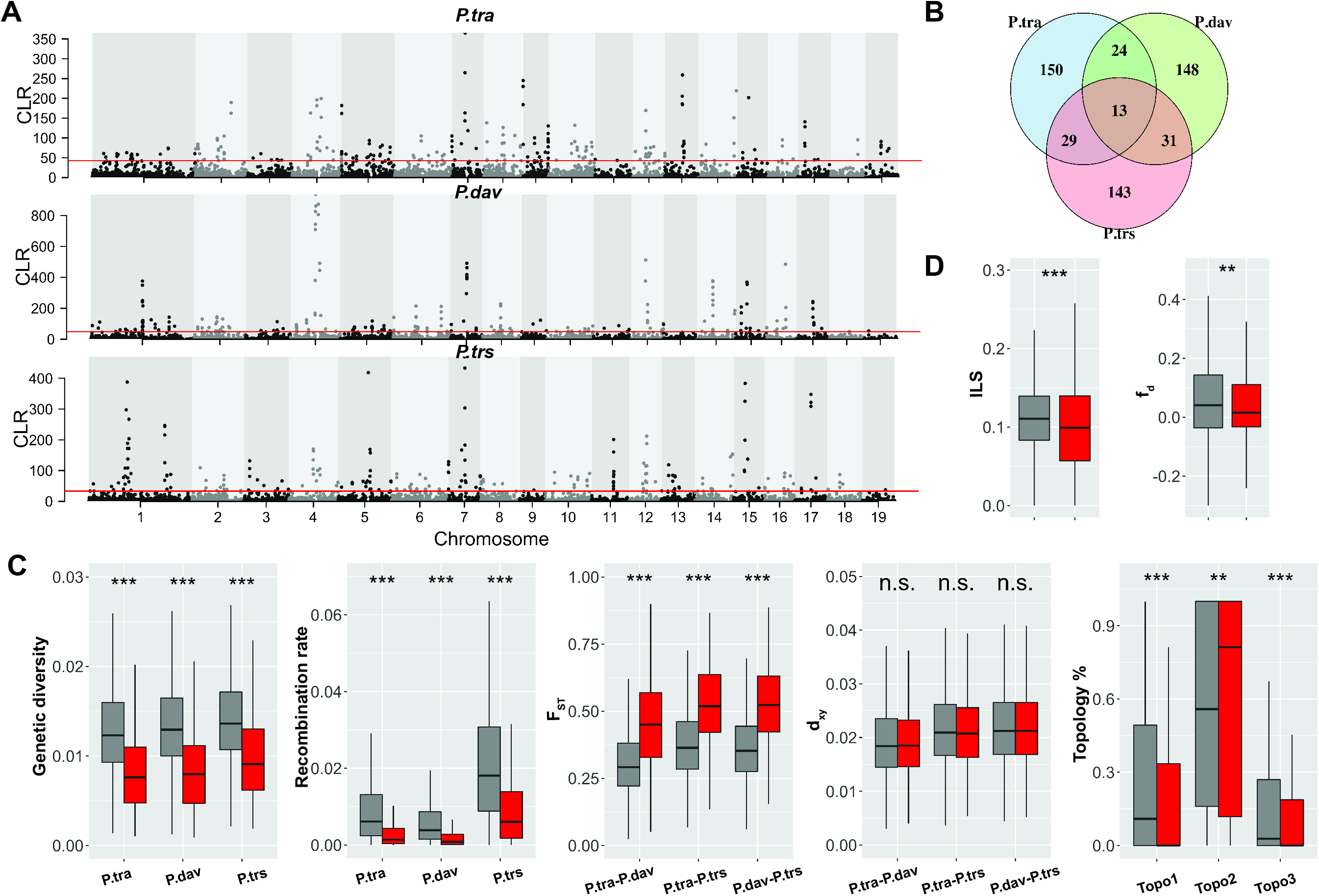
Identification of positive selection. (A) Positive selection analysis by SweepFinder2 reveals windows that are candidates for being under positive selection in the three aspen species: *P. tremula* (*P. tra*), *P. davidiana* (*P. dav*) and *P. tremuloides* (*P. trs*). Horizonal red line indicates the cut-off of composite likelihood ratio (CLR) statistics. (B) The Venn diagram represents shared and unique selected windows detected in the three species. (C) Comparison of genetic diversity, recombination rate, *F*_ST_, d_xy_, and average weightings of the ‘species’ topology between candidate regions under positive selection (red boxes) and genomic background (grey boxes). (D) Comparison of incomplete lineage sorting (ILS) and the estimated admixture proportion (*f*_d_)) between candidate regions under positive selection (red boxes) and genomic background (grey boxes). Asterisks designate significant differences between candidate positive selected regions and the rest of genomic regions by Mann-Whitney *U*-test (^n.s.^ Not significant; * *P* value<0.01; ** *P* value<0.001; *** *P* value <1e-4).

To further study how long-term balancing selection may have driven the evolution of the genomic landscape during speciation, we used a summary statistics, β (beta score), to search for signals of balancing selection across the genome for each aspen species (Siewert & Voight, 2017). As we did for positive selection, we only consider variants with β scores falling in the top 1% as candidate variants. Furthermore, variants simultaneously detected in all three species are considered as potential targets of long-term balancing selection. With this criteria we identified a total of 519 variants putatively under long-term balancing selection across the three aspen species (Figure 6A,B). These variants were unevenly distributed in the genome, and to make them comparable to our previous analyses we clustered them into 32 regions of 10 Kbp windows (Figure 6A). We found significantly higher nucleotide diversity, higher recombination rates, lower *F*_ST_, and higher d_xy_ in the regions under balancing selection compared to the genomic background (Figure 6C). Moreover, we found lower weightings of the ‘species topology’, higher ILS, and lower *f*_d_ in the candidate balancing selection regions although the results were not significant, likely due to the small number of windows showing evidence for balancing selection (Figure 6C,D). We therefore infer that long-term balancing selection may not only influence the genomic landscape of diversity and divergence but may also play a role in shaping the genealogical relationship and barriers to introgression among closely related species (Charlesworth, 2006; Wang et al., 2019).

**Figure 6.**
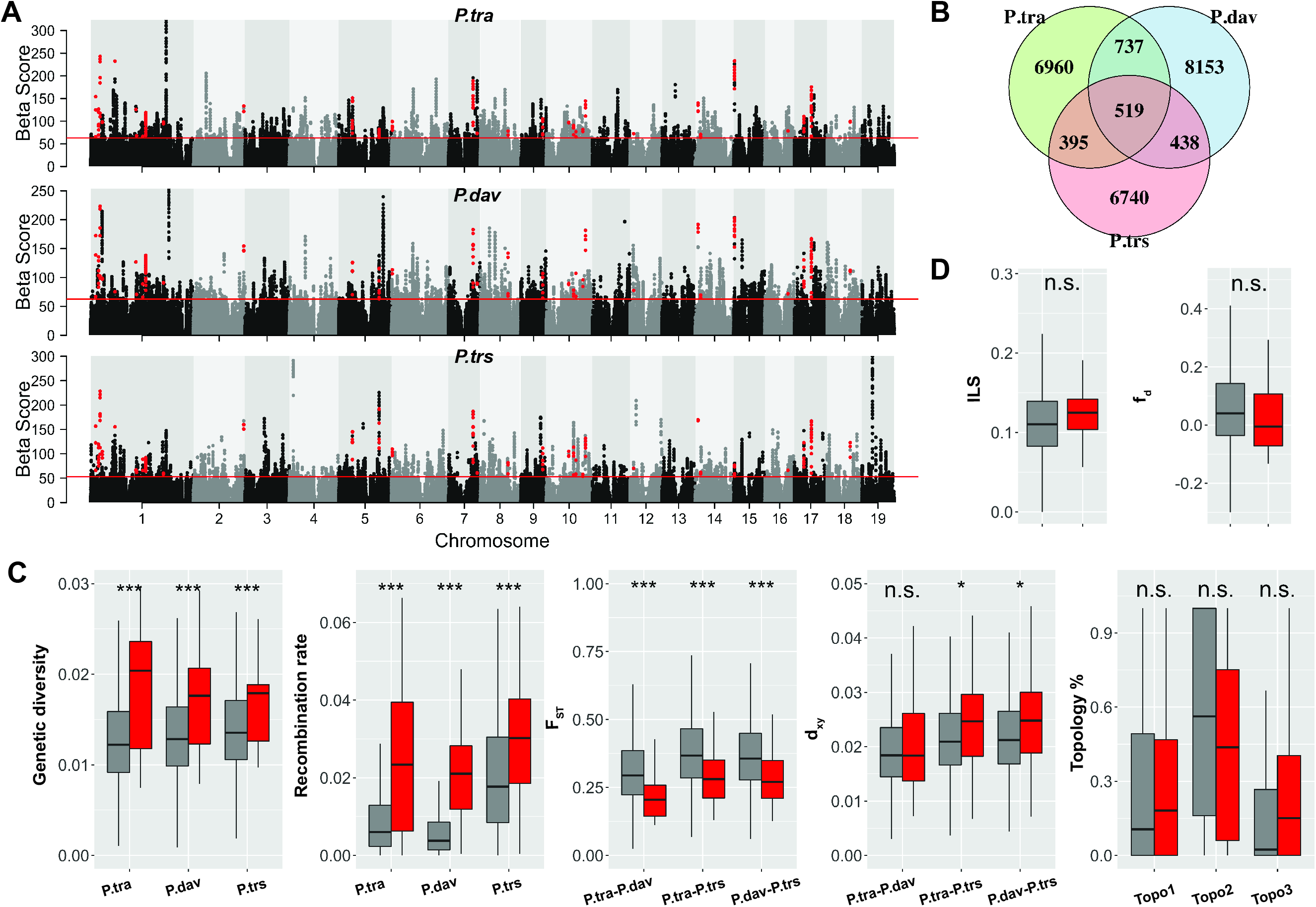
Identification of long-term balancing selection. (A) Signals of balancing selection across all chromosomes in the three aspen species: *P. tremula* (*P. tra*), *P. davidiana* (*P. dav*) and *P. tremuloides* (*P. trs*). Horizonal red line indicates the cut-off of the β statistics. Only the signals detected in all three aspen species (red dots) were considered as being under long-term balancing selection. (B) The Venn diagram represents shared and unique selected SNPs detected in the three species. (C) Comparison of genetic diversity, recombination rate, *F*_ST_, d_xy_, and average weightings of the ‘species’ topology between candidate regions under long-term balancing selection (red boxes) and genomic background (grey boxes). (D) Comparison of incomplete lineage sorting (ILS) and the estimated admixture proportion (*f*_d_)) between candidate regions under long-term balancing selection (red boxes) and genomic background (grey boxes). Asterisks designate significant differences between candidate balancing selected regions and the rest of genomic regions by Mann-Whitney *U*-test (^n.s.^ Not significant; * *P* value<0.01; ** *P* value<0.001; *** *P* value <1e-4).

Finally, to assess whether there were any specific biological functions that were significantly over-represented on genes located in regions identified as undergoing either positive (506 genes) or long-term balancing selection (32 genes), we performed gene ontology (GO) enrichment analysis. We did not detect over-representation for any functional category among the candidate genes under long-term balancing selection. In contrast, we identified 31 significantly enriched GO categories (Fisher’s exact test, *P*<0.01) for genes under positive selection (Table S23). These GO clusters were primarily associated with metabolic processes (DNA, nucleic acid, cellular macromolecule and aromatic compound, molybdopterin confactor), biosynthetic processes (molybdopterin confactor, vitamin B6), cell morphogenesis and gene expression regulation. Together these functional clusters are biologically relevant for plant adaptation, because the biosynthesis of a panoply of diverse natural chemicals serve as important adaptive strategies for sessile long-lived trees to adapt to ever-changing abiotic and biotic environments (Weng, 2014).

## 4. Discussion

Much of our knowledge of how genomic landscape builds in the speciation process is drawn from studies focusing on two young species pairs with ongoing gene flow. Very few examples of now-allopatric species pairs along the speciation continuum have been investigated. Here, we focus our research on four widespread *Populus* species that are allopatrically distributed in northern Hemisphere. After characterizing their speciation and demographic histories, we find that species in different continent exhibited idiosyncratic responses to Pleistocene climate changes. In addition, ancient gene flow was detected between extant Eurasian and North American aspen species (*P. tremula* and *P. tremuloides*). We also investigated the evolutionary forces that have shaped genome-wide patterns of variations within and between species. Our results have found substantial variation in genetic diversity, divergence, species relationships and the extent of introgression along the genome. Variation in these patterns is predictable and can be largely explained by genome-wide variation in the strength and extent of both recent and ancient selection, which depends on the recombination rate and the local density of functional sites. Our findings therefore provide evidence of how recurrent selection interacts with genomic features to shape the genomic landscape during species divergence. We further demonstrate that not only background selection, positive and long-term balancing selection also play crucial roles in shaping genomic variation and phylogenetic relationship among the recently diverged aspen species. Overall, this study highlights the striking impacts of natural selection in shaping within- and between-species genomic variation through speciation.

## Acknowlegements

All analyses were performed on resources provided by the Swedish National Infrastructure for Computing (SNIC) at Uppsala Multidisciplinary Center for Advanced Computational Science (UPPMAX) under the projects SNIC2016-7-89 and SNIC 2017/1-499. Financial support was provided by National Natural Science Foundation of China (31971567) and the Fundamental Research Funds for the Central Universities.

## Data Accessibility Statement

Raw whole genome resequencing data generated for this study have been deposited in the NCBI short read archive under accession number PRJNA576115

## Author Contributions

J.W. conceived the study, analyzed the data and wrote the manuscript. E.J.P. provided the materials of P. davidiana used in this study. N.R.S., J.L., P.K.I. read and commented on the manuscript. All authors approved the final manuscript.

## Notes

#### Summary of Updates

upload vector plots

