## Supplementary Figures for "Evidence for widespread selection in shaping the genomic landscape during speciation of *Populus*"

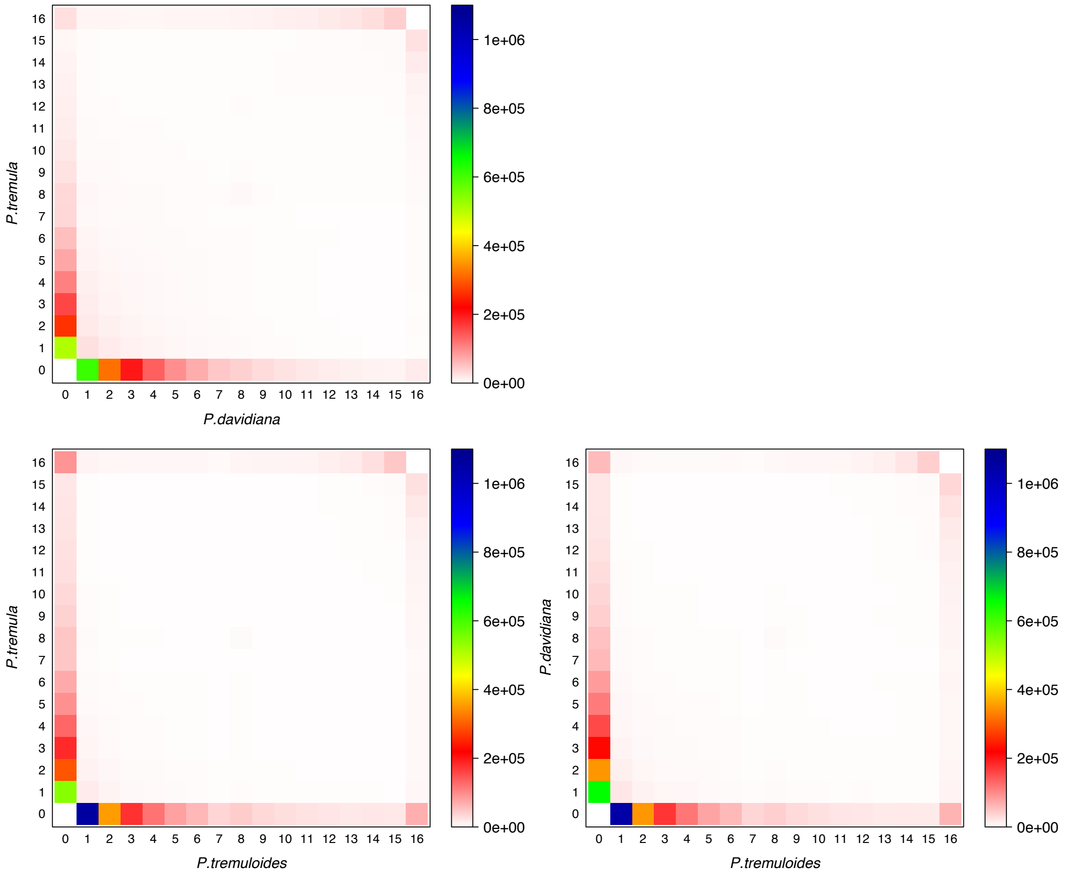


**Figure S1.** Joint allele frequency spectra between *P. tremula*, P. *davidiana* and *P. tremuloides*. Colors reflect the number of SNPs.

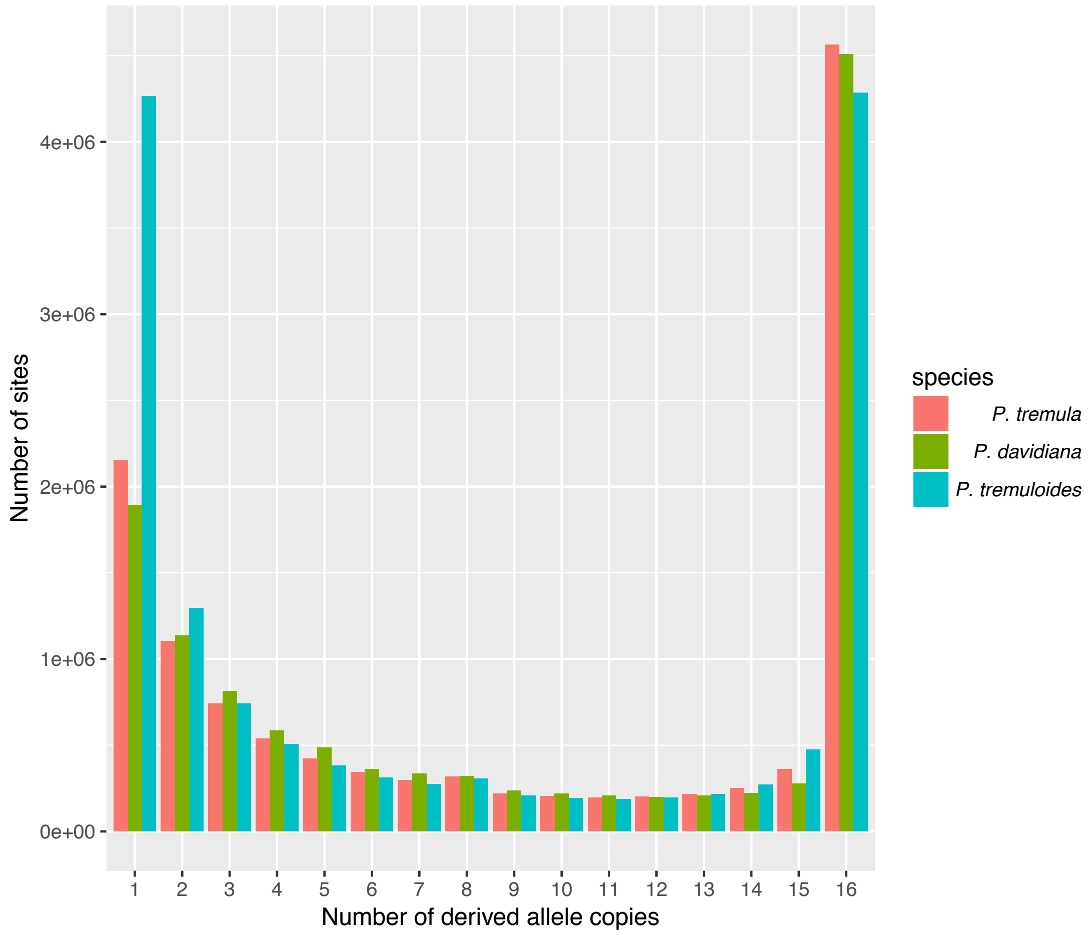


**Figure S2.** Comparison of the derived site frequency spectrum of three aspen species: *P. tremula*, *P. davidiana* and *P. tremuloides*.

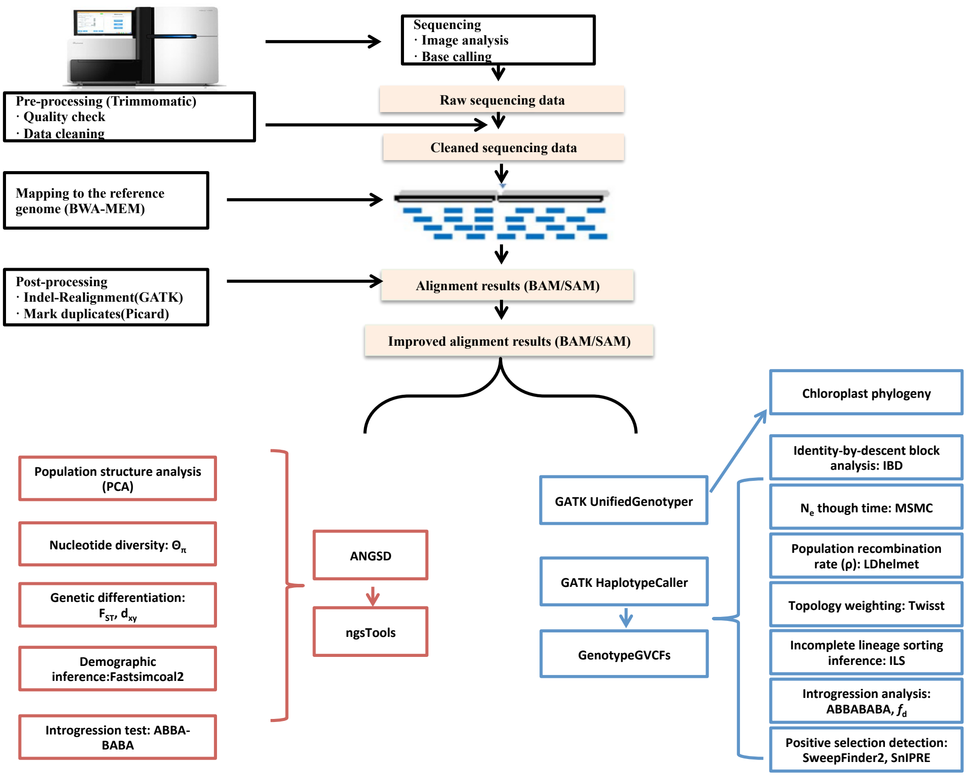


**Figure S3.** Analysis workflow used in this study.

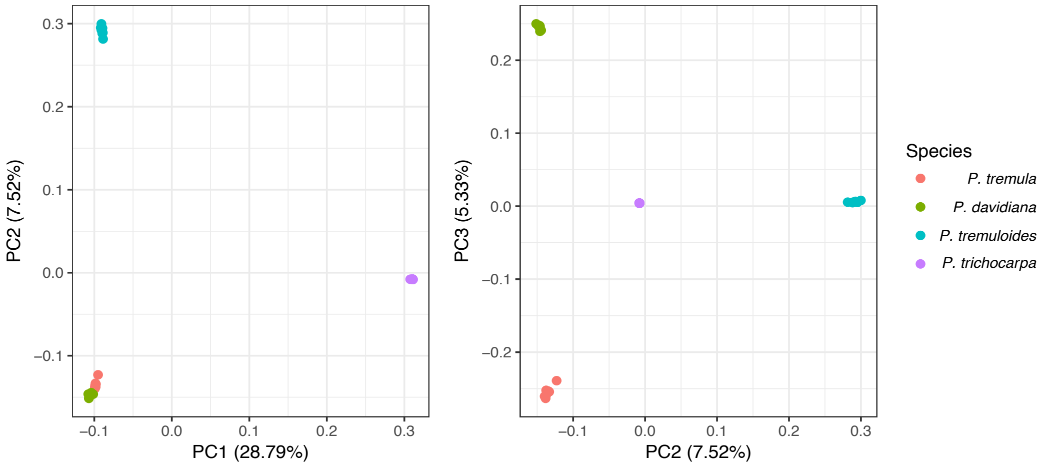


**Figure S4.** Principal component analysis (PCA) based on genetic covariance among all individuals of the four *Populus* species. Percent variation explained by each component is shown in parentheses.

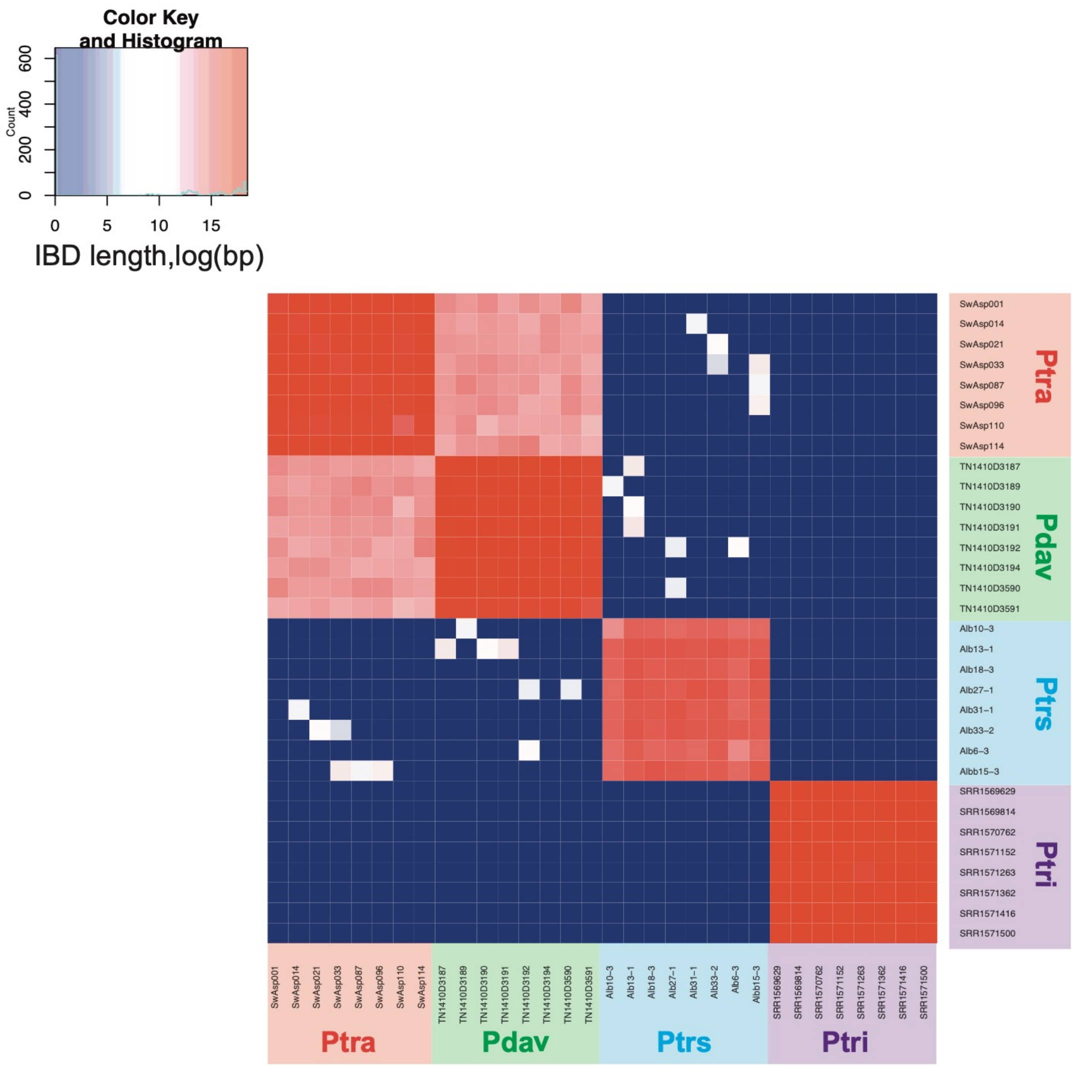


**Figure S5**. Estimated haplotype sharing between individuals of four *Populus* species: *P. tremula* (Ptra), *P. davidiana* (Pdav), *P. tremuloides* (Ptrs) and *P. trichocarpa* (Ptri). Heatmap colors represent the total length of identical-by-descent (IBD) blocks for each pairwise comparison.

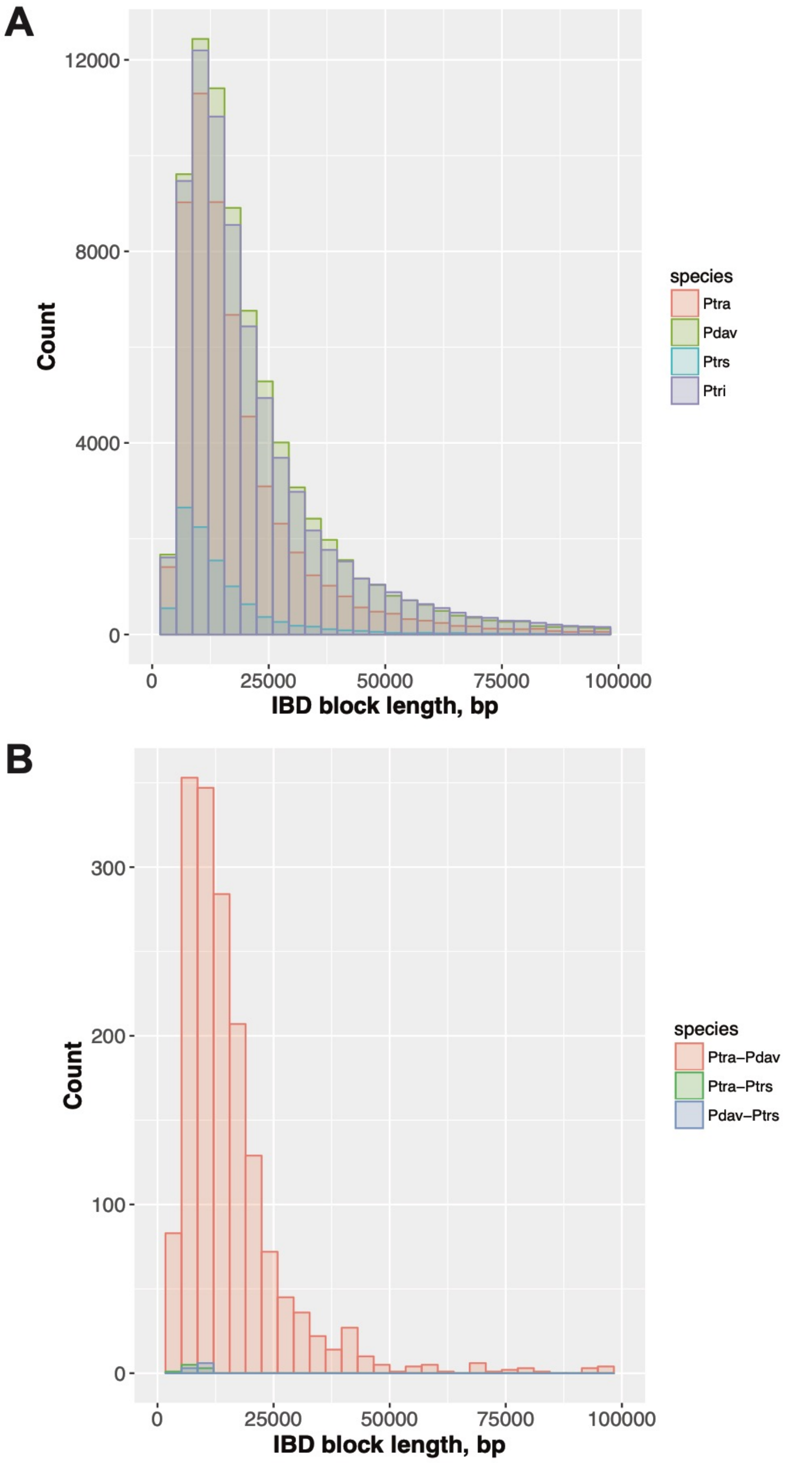


**Figure S6.** (A) Distribution of shared identical-by-descent (IBD) blocks length identified within species: *P. tremula* (Ptra), *P. davidiana* (Pdav), *P. tremuloides* (Ptrs) and *P. trichocarpa* (Ptri). (B) Distribution of shared IBD blocks length between pairs of the three aspen species.

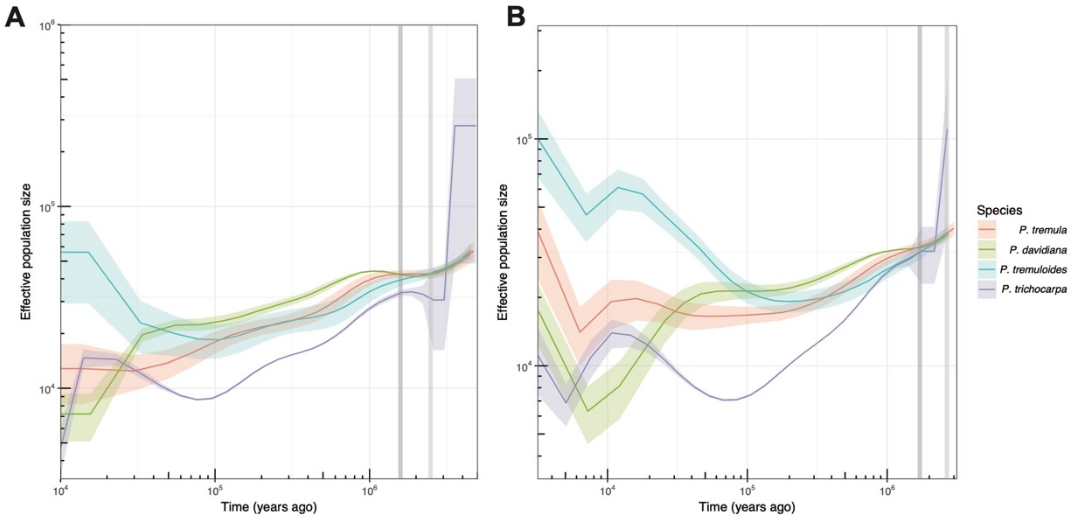


**Figure S7.** Historical effective population size of the four *Populus* species inferred using MSMC v2 based on sets of two haplotypes (A) and four haplotypes (B), respectively. Solid lines represent medians and shading represents ± standard deviation calculated across pairs of haplotypes.

**model1**

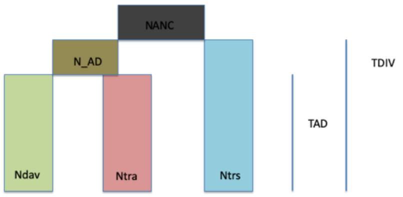


**model2**

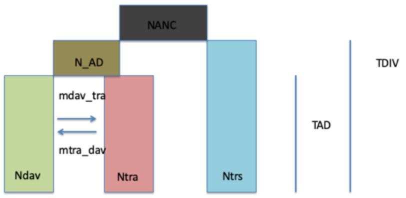


**model3**

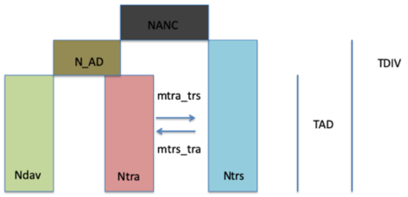


**model4**

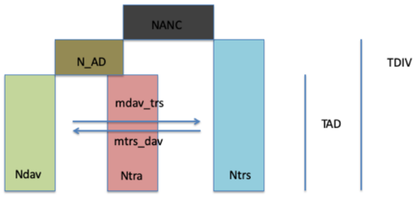


**model5**

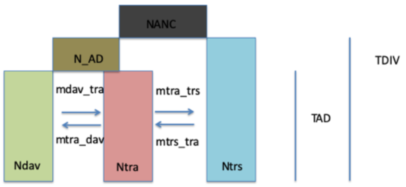


**model6**

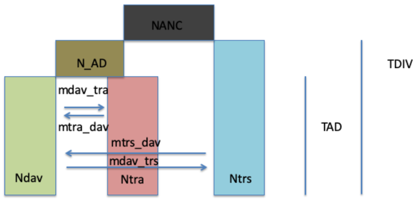


**model7**

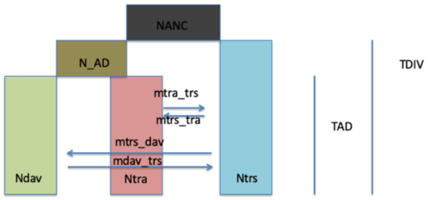


**model8**

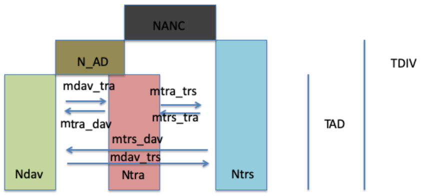

**model9**

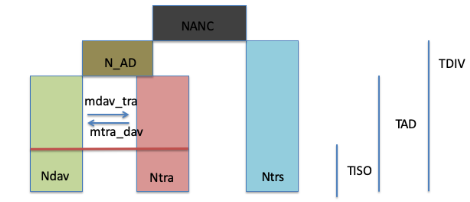


**model10**

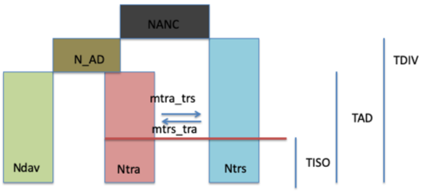


**model11**

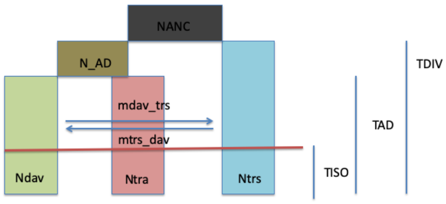


**model12**

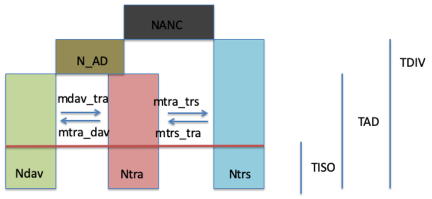


**model13**

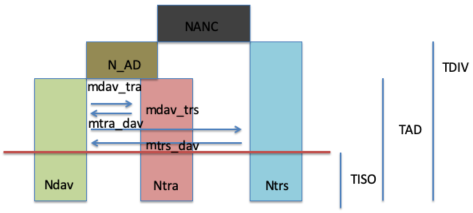


**model14**

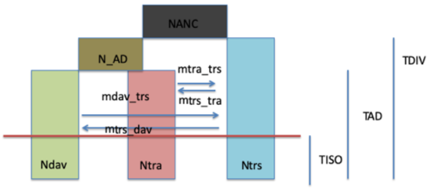


**model15**

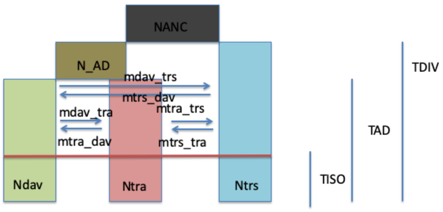


**model16**

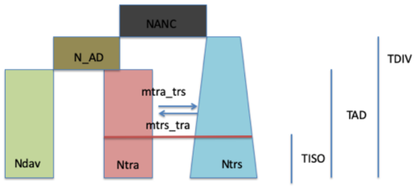

**model17**

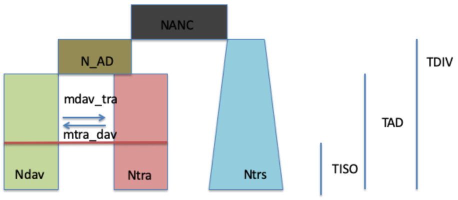


**model18**

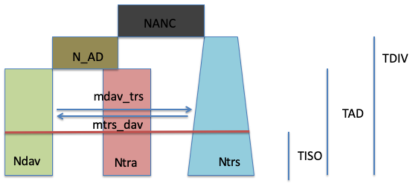


**model19**

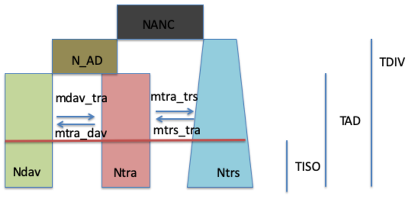


**model20**

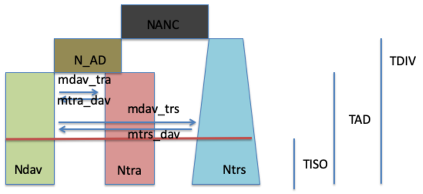


**model21**

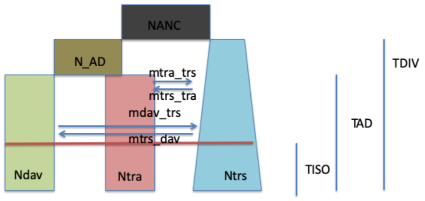


**model22**

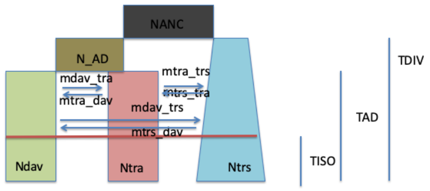


**model23**

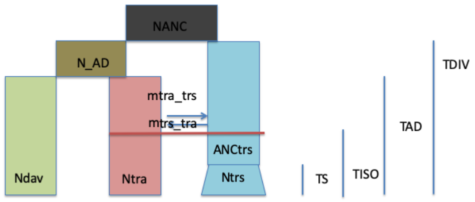


**model24**

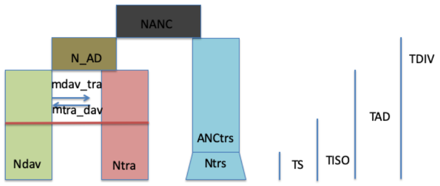

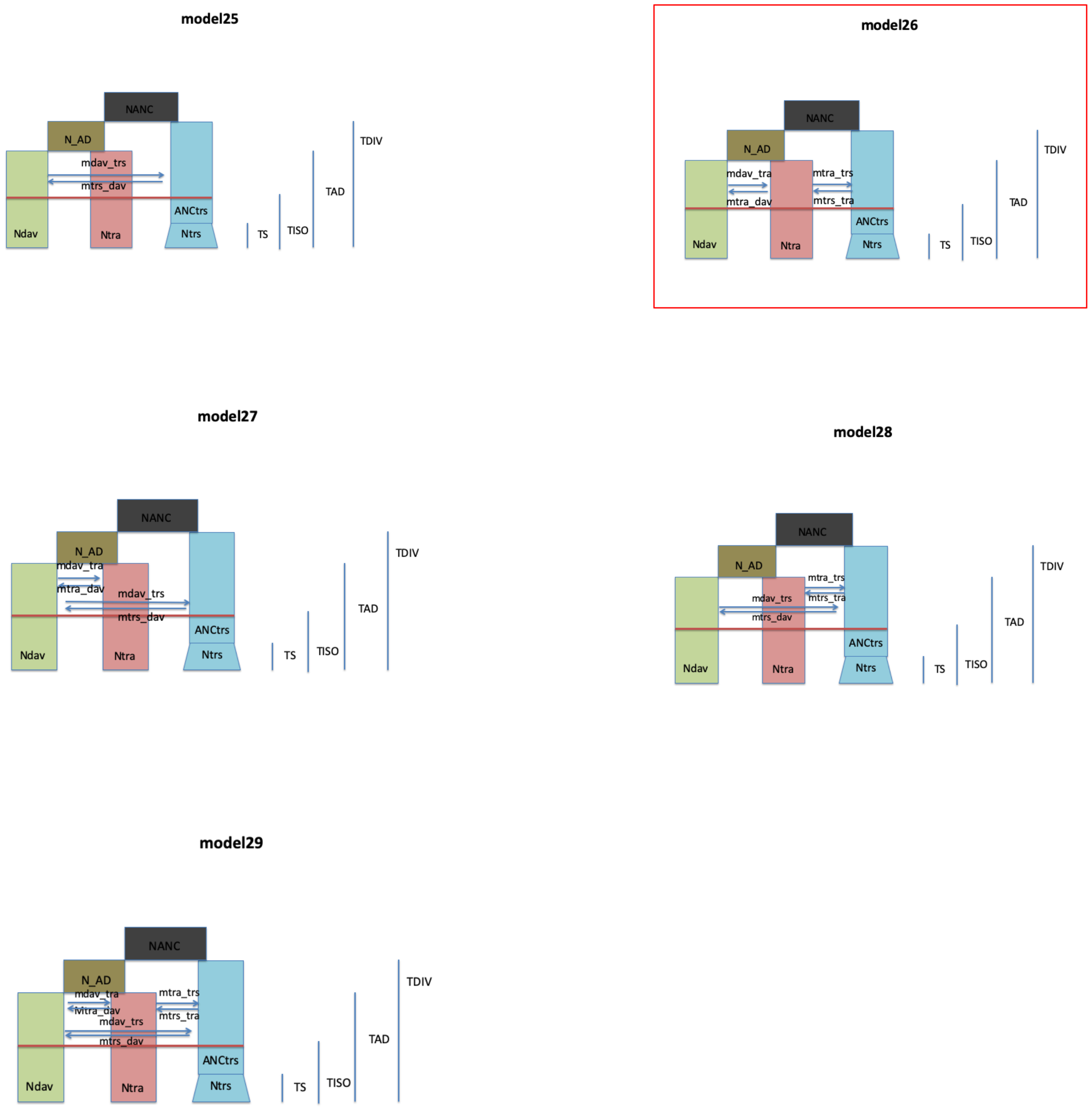


**Figure S8.** Twenty-nine tested demographic models for the speciation process of aspens. Model 1-Model 8, all three aspen species have constant population size after splitting, differed by isolation -with or -without asymmetric gene flow between different pairs of species. Model 9-Model 15, all three aspen species have constant population size after splitting, differed by isolation with ancestral asymmetric gene flow between different pairs of species occurred before time point TISO. Model 16-Model 22, *P. tremula* and *P. davidiana* have constant population size after splitting and *P. tremuloides* experienced exponential population size change after splitting. The models differed by isolation with ancestral asymmetric gene flow between different pairs of species occurred before time point TISO. Model 23-Model 29, *P. tremula* and *P. davidiana* have constant population size after splitting and *P. tremuloides* expanded exponentially at a recent time TS. The models differed by isolation with ancestral asymmetric gene flow occurred between different pairs of species before time point TISO.

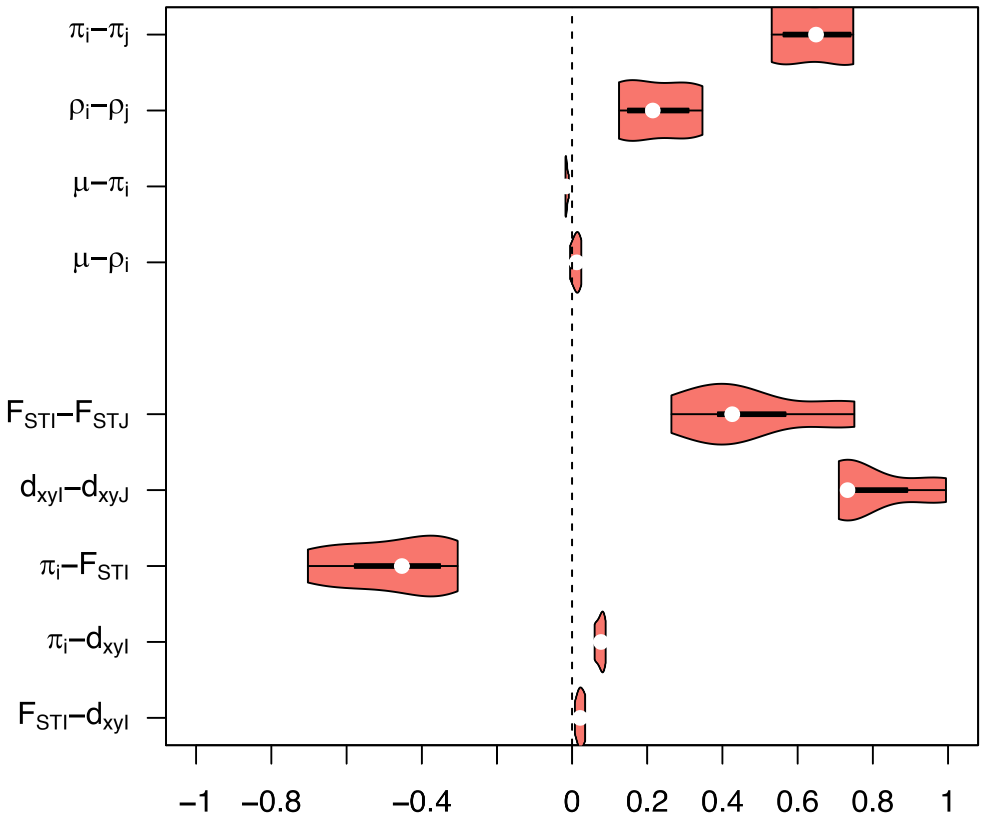


**Figure S9.** Distribution of correlation coefficients (Spearman's ρ) shown as violin plots for population summary statistics characterizing genomic features (neutral mutation rate $\mu$) and variation within ($\pi$, $\rho$) and between species ($F_{ST}$, $d_{xy}$) calculated at 10 Kbp windows. Subscripts 'i, j' symbolize all possible combinations of correlations between two species i=1…(n-1) and j=(i+1)…n for within-species measures; Capital letters 'I, J' symbolize inter-species statistics. Correlations exclude pseudo-replicated species comparisons.

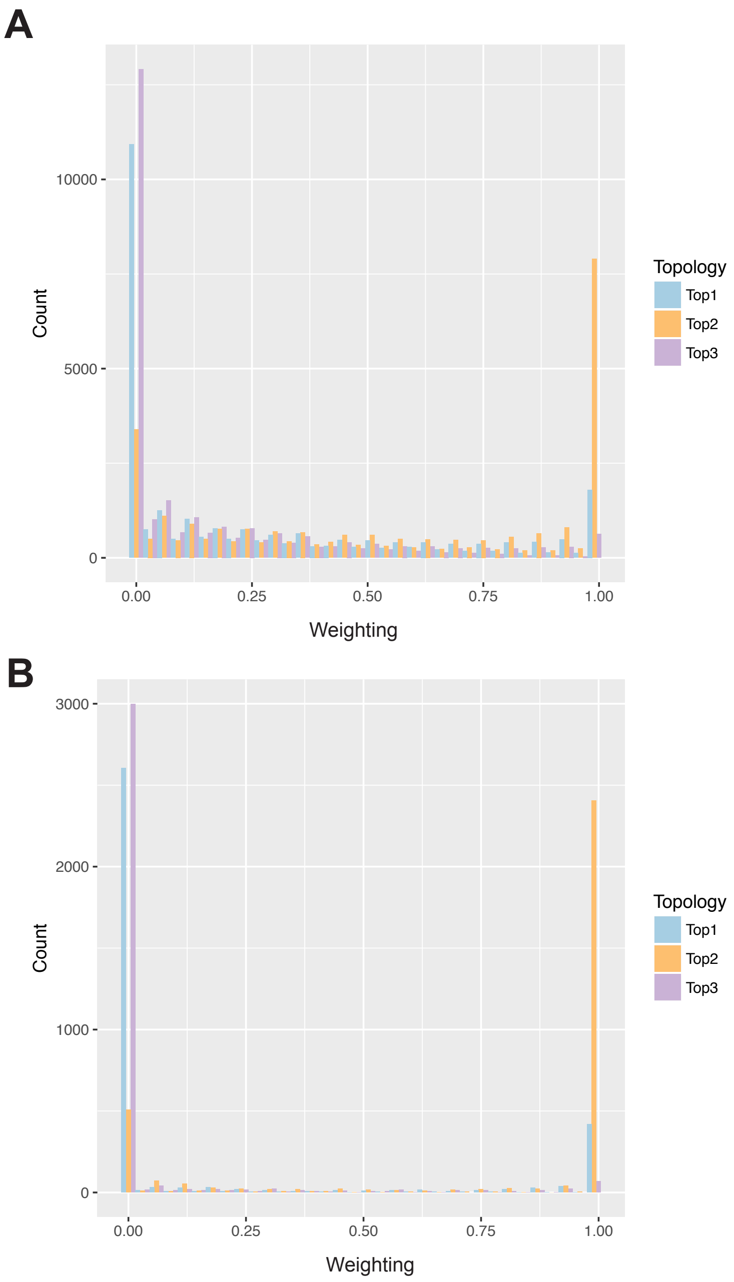


**Figure S10.** Comparison of the distribution of topology weighting for the three topologies of aspen species in (A) 10 Kbp and (B) 100 Kbp windows. Topology (Top)1: [[*P. tremula*, *P. tremuloides*], *P. davidiana*]; Top2: [[*P. tremula*, *P. davidiana*], *P. tremuloides*]; Top3: [[*P. davidiana*, *P. tremuloides*], *P. tremula*]

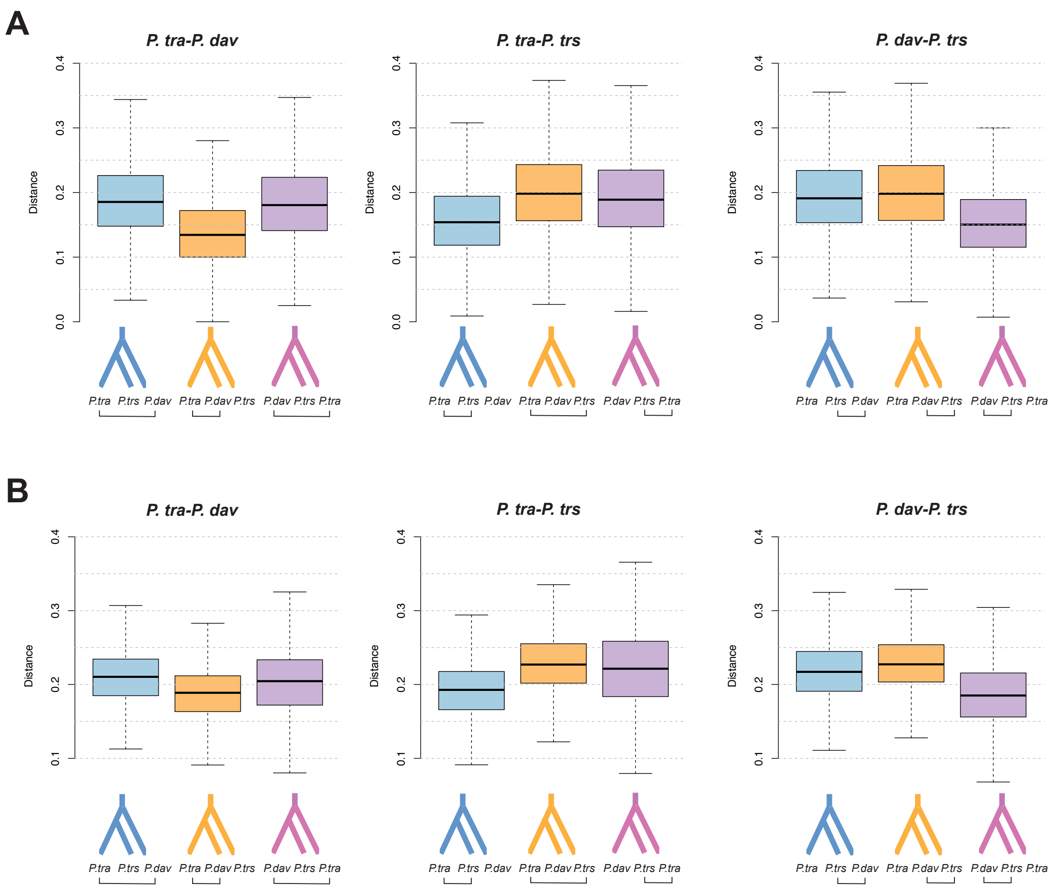


**Figure S11.** Average branch length separating each pair of aspen species, *P. tremula* (*Ptra*), *P. davidiana* (*Pdav*), *P. tremuloides* (*Ptrs*), in different sub-trees in (A) 10 Kbp and (B) 100 Kbp windows. Boxplot show the distribution of average pairwise distance across all trees, separated by the topology matched by each sub-tree (shown below and colored). For example, the blue box in the left-hand plot gives the distribution of average pairwise distances between samples from species *P. tra* and *P. dav* for all sub-trees that matched the topology '[[*P. tra*, *P. dav*], *P. trs*]'. The corresponding distances are plotted for the other two topologies in different colors, and then for the other pairwise comparisons (*P. tra-P. trs* and *P. dav-P. trs*) in the middle and right-hand plots, respectively.

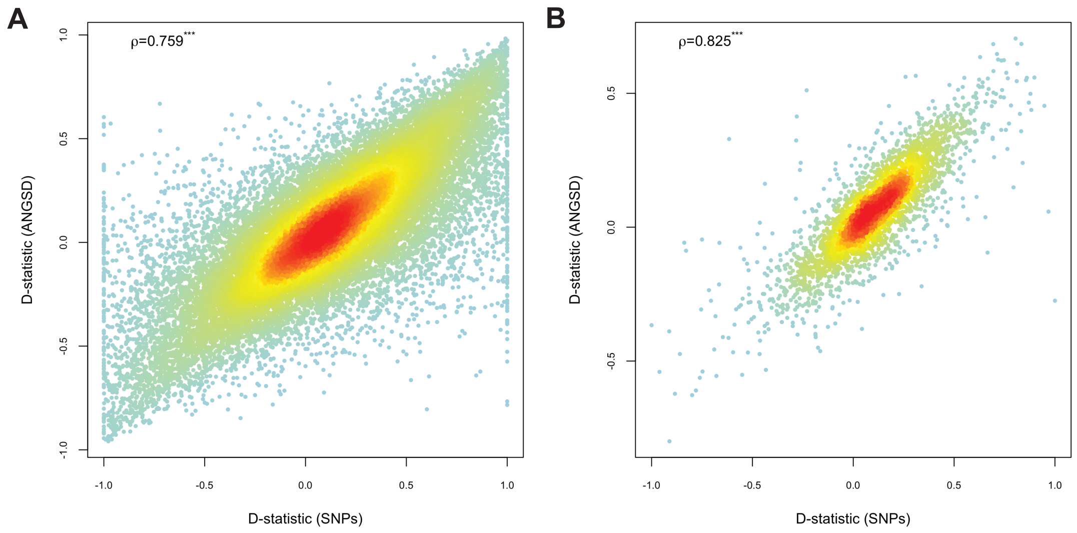


**Figure S12.** Relationship between the *D*-statistic values estimated based on genotype likelihood (ANGSD) and those with called genotypes (SNPs) in (A) 10 Kbp and (B) 100 Kbp non-overlapping windows across the genome. The red to yellow to blue gradient indicates decreased density of observed events at a given location in the graph.

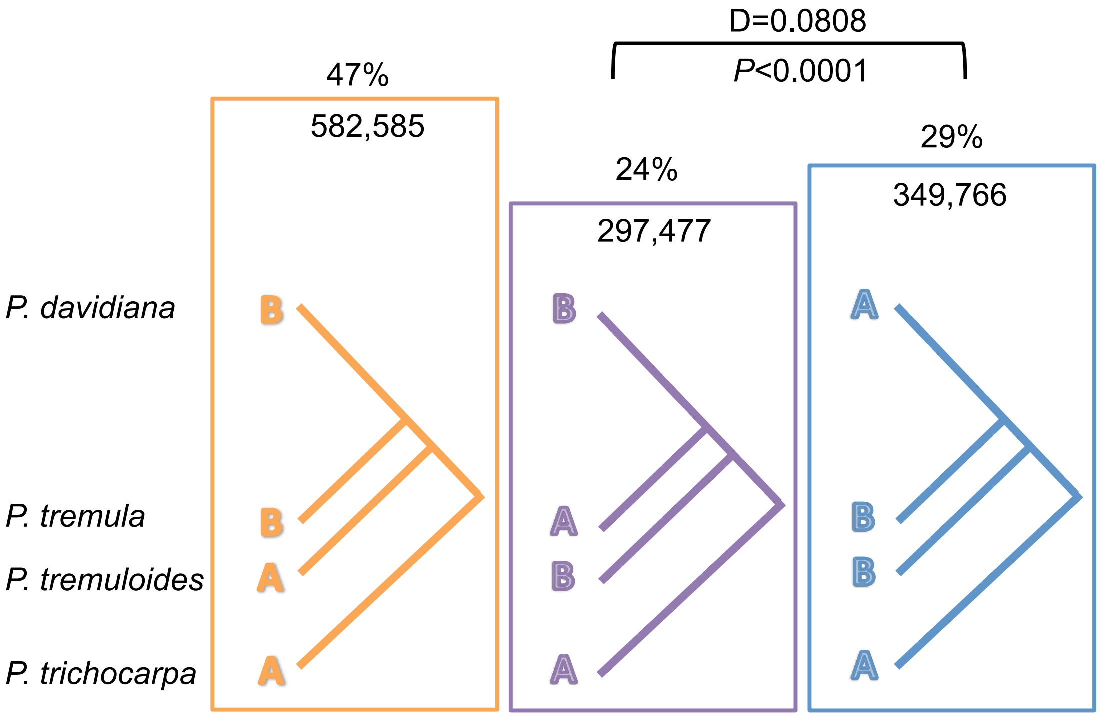


**Figure S13.** ABBA-BABA analysis of *P. tremula*, *P. davidiana*, *P. tremuloides* and *P. trichocarpa*. Number of sites supporting different trees is indicated both as a percentage and as actual numbers. The *D* statistic and corresponding block jackknife corrected *P* value are given for testing the null hypothesis of symmetry in genetic relationships.

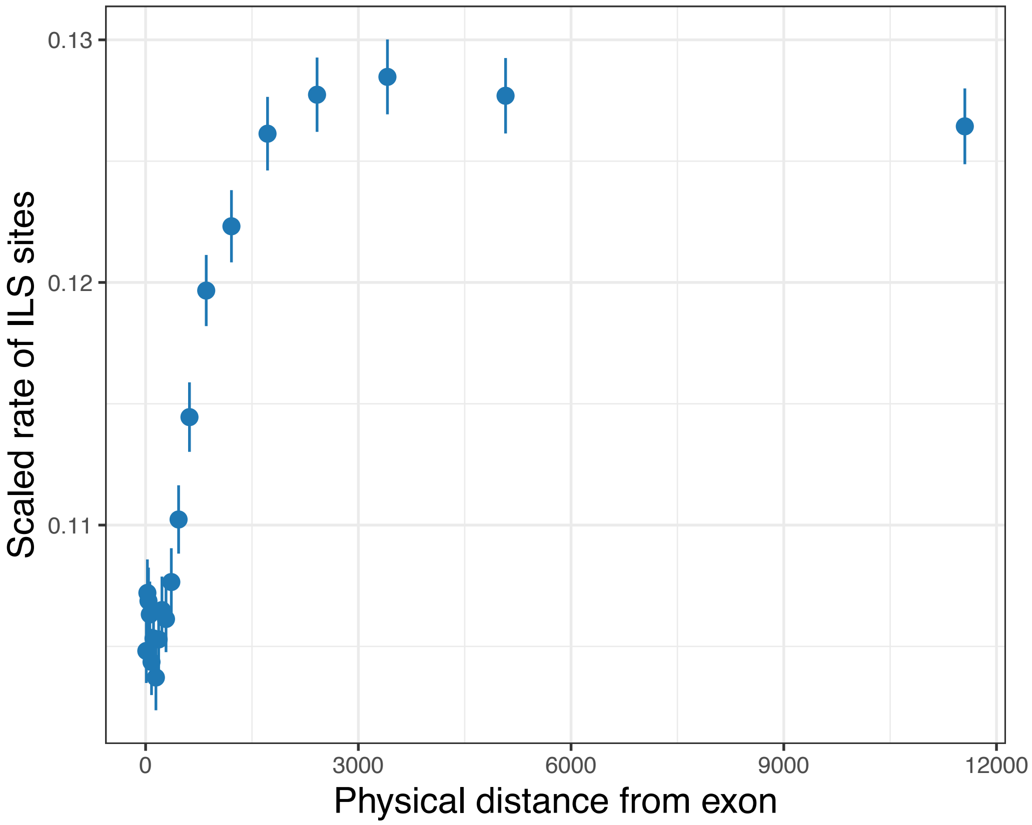


**Figure S14.** Reduction in incomplete lineage sorting (ILS) as a function of physical distance to the nearest exon. Bins are selected to put approximately the same amount of sites in each bin. Error bars show 1.96×standard error.

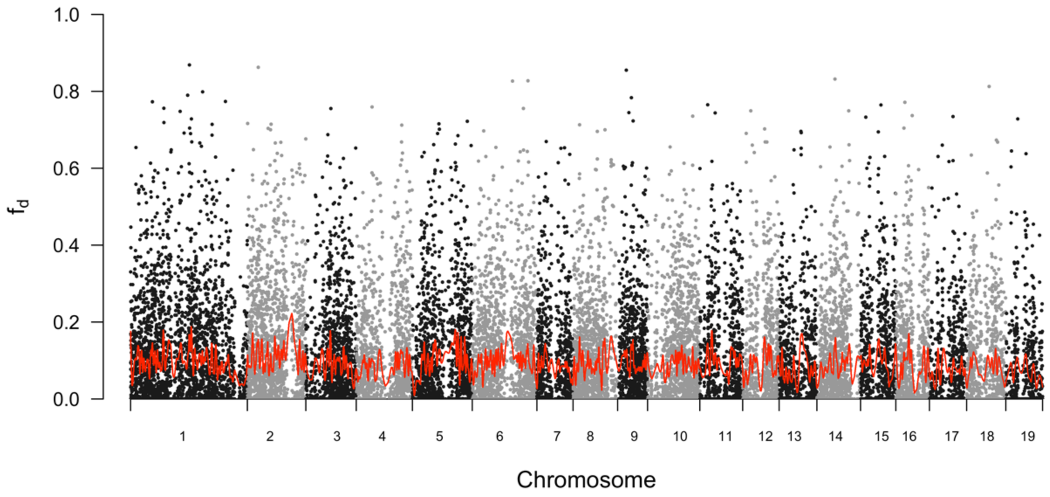


**Figure S15.** Estimated admixture proportion ($f_d$) between *P. tremula* and *P. tremuloides* plotted across all 19 chromosomes in 10 Kbp non-overlapping windows (grey and black dots), with locally weighted averages being plotted as red lines.
