## Supplementary Tables for "Evidence for widespread selection in shaping the genomic landscape during speciation of *Populus*"

**Table S1.** Samples in this study

| SampleID | Site | Latitude | Longitude | Coverage | SRA |
|---|---|---|---|---|---|
| ***P. tremula*** | | | | | |
| SwAsp001 | Simlang | 56.6925 | 13.2147 | 20.50 | SRR2744682 |
| SwAsp014 | Ronneby | 56.3081 | 15.1269 | 24.53 | SRR2745905 |
| SwAsp021 | Vargarda | 57.9917 | 12.9119 | 18.50 | SRR2745906 |
| SwAsp033 | Ydre | 57.8281 | 15.3103 | 20.11 | SRR2745909 |
| SwAsp087 | Dorotea | 64.3406 | 16.3992 | 24.44 | SRR2746774 |
| SwAsp096 | Umea | 63.9781 | 20.7056 | 23.43 | SRR2746685 |
| SwAsp110 | Arjeplog | 66.2592 | 18 | 36.83 | SRR2746848 |
| SwAsp114 | Lulea | 65.5544 | 22.3939 | 34.62 | SRR2746922 |
| ***P. davidiana*** | | | | | |
| Daehwa6-1 | Taehwa | 37.5018 | 128.4582 | 22.51 | SAMN12924941 |
| Daehwa18-2 | Taehwa | 37.5018 | 128.4582 | 22.62 | SAMN12924942 |
| Palgong1-1 | Palkong | 36.0167 | 128.6903 | 23.98 | SAMN12924943 |
| Palgong3-1 | Palkong | 36.0167 | 128.6903 | 23.39 | SAMN12924944 |
| Palgong2-3 | Palkong | 36.0167 | 128.6903 | 28.80 | SAMN12924945 |
| Bonghyeon4_2 | Ponghyun | 37.8262 | 128.4868 | 24.08 | SAMN12924946 |
| Dongdu2-1 | Wondoo | 37.7042 | 128.44 | 23.68 | SAMN12924947 |
| Sogwang9-1 | Sokwang | 36.9927 | 128.1917 | 29.47 | SAMN12924948 |
| ***P. tremuloides*** | | | | | |
| Alb10-3 | Alberta | 51.0718 | -115.0044 | 21.08 | SRR2748652 |
| Alb13-1 | Alberta | 51.0479 | -115.0232 | 23.22 | SRR2748654 |
| Alb18-3 | Alberta | 51.0686 | -115.3516 | 21.49 | SRR2748658 |
| Alb27-1 | Alberta | 51.0405 | -114.8939 | 29.02 | SRR2748660 |
| Alb31-1 | Alberta | 51.0435 | -114.8352 | 23.06 | SRR2748661 |
| Alb33-2 | Alberta | 51.0431 | -114.7568 | 27.76 | SRR2748662 |
| Alb6-3 | Alberta | 51.1324 | -115.0664 | 28.77 | SRR2749823 |
| Albb15-3 | Alberta | 51.0811 | -115.3767 | 22.14 | SRR2749863 |
| ***P. trichocarpa*** | | | | | |
| BESC-56 | Talley_Way | 46.099 | -122.878 | 25.43 | SRR1571263 |
| BESC-281 | Monroe | 47.851 | -121.962 | 22.89 | SRR1571362 |
| BESC-840 | Orting | 47.042 | -122.209 | 27.56 | SRR1571416 |
| BESC-873 | Sultan | 47.856 | -121.811 | 24.36 | SRR1571152 |
| DEND-17-2 | DEND | 52.817 | -126.95 | 15.26 | SRR1569629 |
| GW-9598 | Nisqually_River | 47.067 | -123.733 | 21.84 | SRR1571500 |
| LILC-26-3 | Harrison | 51.233 | -124.95 | 28.53 | SRR1569814 |
| SLMB-28-4 | Salmon | 50.217 | -125.817 | 22.33 | SRR1570762 |

**Table S2.** Tracy-Widom statistics for the first three eigenvalues in PCA analysis

| Eigenvectors | Eigenvalues | Twstat | *P*-value |
|---|---|---|---|
| 1 | 28.79 | 6.501 | 5.25329e-07 |
| 2 | 7.52 | 10.642 | 6.93175e-12 |
| 3 | 5.33 | 11.724 | 1.82012e-13 |

**Table S3.** The average number of identical-by-descent (IBD) haplotypes within and between four *Populus* species

| Number | *P. tremula* | *P. davidiana* | *P. tremuloides* | *P. trichocarpa* |
|---|---|---|---|---|
| *P. tremula* | 56487 | - | - | - |
| *P. davidiana* | 1684 | 78090 | - | - |
| *P. tremuloides* | 9 | 9 | 10378 | - |
| *P. trichocarpa* | 0 | 0 | 0 | 76221 |

**Table S4.** The average length of identical-by-descent (IBD) haplotypes within and between four *Populus* species

| Number | *P. tremula* | *P. davidiana* | *P. tremuloides* | *P. trichocarpa* |
|---|---|---|---|---|
| *P. tremula* | 21687 | - | - | - |
| *P. davidiana* | 18028 | 25709 | - | - |
| *P. tremuloides* | 7616 | 9691 | 18744 | - |
| *P. trichocarpa* | 0 | 0 | 0 | 27624 |

**Table S5.** Statistics of nucleotide diversity (π) (mean ± standard error) over 10 Kbp non-overlapping windows along 19 chromosomes for the four *Populus* species

| Chromosome | *P. tremula* | *P. davidiana* | *P. tremuloides* | *P. trichocarpa* |
|---|---|---|---|---|
| Chr01 | 0.0145±0.0001 | 0.0150±0.0001 | 0.0156±0.0001 | 0.0066±0.0001 |
| Chr02 | 0.0128±0.0001 | 0.0136±0.0001 | 0.0140±0.0001 | 0.0057±0.0001 |
| Chr03 | 0.0137±0.0002 | 0.0143±0.0002 | 0.0148±0.0002 | 0.0061±0.0001 |
| Chr04 | 0.0137±0.0002 | 0.0138±0.0002 | 0.0152±0.0002 | 0.0064±0.0002 |
| Chr05 | 0.0127±0.0001 | 0.0130±0.0001 | 0.0141±0.0001 | 0.0060±0.0001 |
| Chr06 | 0.0133±0.0001 | 0.0140±0.0001 | 0.0145±0.0001 | 0.0058±0.0001 |
| Chr07 | 0.0133±0.0002 | 0.0135±0.0002 | 0.0143±0.0002 | 0.0065±0.0001 |
| Chr08 | 0.0116±0.0001 | 0.0123±0.0001 | 0.0130±0.0001 | 0.0052±0.0001 |
| Chr09 | 0.0119±0.0002 | 0.0124±0.0002 | 0.0136±0.0002 | 0.0052±0.0001 |
| Chr10 | 0.0119±0.0001 | 0.0125±0.0001 | 0.0133±0.0001 | 0.0055±0.0001 |
| Chr11 | 0.0153±0.0002 | 0.0158±0.0002 | 0.0166±0.0002 | 0.0073±0.0002 |
| Chr12 | 0.0143±0.0002 | 0.0146±0.0002 | 0.0158±0.0002 | 0.0066±0.0002 |
| Chr13 | 0.0137±0.0002 | 0.0144±0.0002 | 0.0145±0.0002 | 0.0064±0.0002 |
| Chr14 | 0.0129±0.0002 | 0.0131±0.0002 | 0.0144±0.0002 | 0.0060±0.0001 |
| Chr15 | 0.0145±0.0002 | 0.0146±0.0002 | 0.0154±0.0002 | 0.0071±0.0002 |
| Chr16 | 0.0152±0.0002 | 0.0158±0.0002 | 0.0164±0.0002 | 0.0071±0.0001 |
| Chr17 | 0.0154±0.0003 | 0.0156±0.0002 | 0.0163±0.0002 | 0.0077±0.0002 |
| Chr18 | 0.0140±0.0002 | 0.0147±0.0002 | 0.0153±0.0002 | 0.0070±0.0002 |
| Chr19 | 0.0161±0.0003 | 0.0169±0.0003 | 0.0178±0.0003 | 0.0087±0.0002 |
| Genomewide | 0.0135±0.0000 | 0.0141±0.0000 | 0.0148±0.0000 | 0.0063±0.0000 |

**Table S6**. Relative likelihood of the different models shown in Figure S8.

| Model | Max(log$_{10}$(Lhoodi)[a] | No. Of parameters(d) | AIC$_i$[b] | $\Delta_i$[b] | Model normalized relative likelihood ($w_i$)[b] |
|---|---|---|---|---|---|
| model1 | -167511807 | 7 | 771420393.4 | 24607289.21 | 0 |
| model2 | -167454404 | 9 | 771156046.8 | 24342942.63 | 0 |
| model3 | -166594146 | 9 | 767194412.3 | 20381308.14 | 0 |
| model4 | -166624079 | 9 | 767332258.9 | 20519154.69 | 0 |
| model5 | -165731928 | 11 | 763223755.7 | 16410651.51 | 0 |
| model6 | -165990143 | 11 | 764412879.7 | 17599775.53 | 0 |
| model7 | -167243451 | 11 | 770184576.3 | 23371472.16 | 0 |
| model8 | -166723580 | 13 | 767790485.9 | 20977381.73 | 0 |
| model9 | -167479611 | 10 | 771272131.3 | 24459027.15 | 0 |
| model10 | -163680022 | 10 | 753774377.4 | 6961273.17 | 0 |
| model11 | -164080374 | 10 | 755618066.5 | 8804962.27 | 0 |
| model12 | -165585240 | 12 | 762548234.5 | 15735130.30 | 0 |
| model13 | -165414901 | 12 | 761763794.4 | 14950690.22 | 0 |
| model14 | -165625338 | 12 | 762732892.6 | 15919788.42 | 0 |
| model15 | -165374586 | 14 | 761578141 | 14765036.78 | 0 |
| model16 | -164050832 | 10 | 755482020.5 | 8668916.33 | 0 |
| model17 | -167906882 | 10 | 773239787 | 26426682.82 | 0 |
| model18 | -163899648 | 10 | 754785792.5 | 7972688.28 | 0 |
| model19 | -165845930 | 12 | 763748756.3 | 16935652.12 | 0 |
| model20 | -163122406 | 12 | 751206464.8 | 4393360.59 | 0 |
| model21 | -163332093 | 12 | 752172109.1 | 5359004.91 | 0 |
| model22 | -162699522 | 14 | 749259016 | 2445911.80 | 0 |
| model23 | -162809774 | 12 | 749766741.2 | 2953637.03 | 0 |
| model24 | -166471569 | 12 | 766629930.4 | 19816826.19 | 0 |
| model25 | -162712187 | 12 | 749317336.5 | 2504232.29 | 0 |
| model26 | -162168399 | 14 | 746813104.2 | 0.00 | 1 |
| model27 | -164383763 | 14 | 757015232.4 | 10202128.24 | 0 |
| model28 | -162185808 | 14 | 746893275.6 | 80171.41 | 0 |
| model29 | -164211090 | 16 | 756220047.9 | 9406943.69 | 0 |

**Table S7.** Inferred demographic parameters of the best-fitting demographic model (model 26 from Figure S8) shown in Figure 2B.

| **Parameters** | **Point estimation** | **95%CI[a]** | |
|---|---|---|---|
| | | **Lower bound** | **Upper bound** |
| Ntra | 163300 | 128872 | 229616 |
| Ndav | 164257 | 126859 | 245915 |
| Ntrs | 1483551 | 542417 | 1483522 |
| ANCtrs | 36843 | 27654 | 94638 |
| N_AD | 6966 | 5454 | 23744 |
| NANC | 2870799 | 726275 | 2870799 |
| TS | 771960 | 440280 | 886635 |
| TISO | 847005 | 538815 | 1017660 |
| TAD | 1694310 | 1470150 | 2083140 |
| TDIV | 2436765 | 2083755 | 3245760 |
| mdav-tra | 3.33097e-05 | 5.695731e-06 | 3.54594e-05 |
| mtra-dav | 2.85859e-08 | 3.813319e-09 | 3.572837e-06 |
| mtra-trs | 3.57121e-06 | 5.662713e-07 | 7.372676e-06 |
| mtrs-tra | 6.26018e-05 | 3.35099e-05 | 6.664749e-05 |

Note: Parameters correspond to the model26 from Figure S8. Ntra, Ndav, Ntrs indicate the effective population size of *P. tremula*, *P. davidiana* and P. *tremulodies*, respectively. ANCtrs indicates the effective population size of *P. tremuloides* before population expansion. N_AD indicate the effective population size of the ancestor population before the split of *P. tremula* and *P. davidiana*.

NANC indicate the effective population size of the ancestral population before the split of *P. tremuloides* and the Euasian lineage. TS indicates the estimated start of population expansion of *P. tremuloides*. TISO indicates the estimated time since the end of gene flow between species. TAD indicates the estimated divergence time between *P. tremula* and *P. davidiana*. TDIV indicates the estimated divergence time between *P. tremuloides* and the ancestor population of *P. tremula* and *P. davidiana*. mdav-tra, mtra-dav, mtra-trs, mtrs-tra indicate the per-generation migration rate from *P. davidiana* to *P. tremula*, from *P. tremula* to *P. davidiana*, from *P. tremula* to *P. tremuloides*, from *P. tremuloides* to *P. tremula*, respectively.

[a] Parametric bootstrap estimates obtained by parameter estimation from 100 datasets simulated according to the overall maximum composite likelihood estimates shown in point estimation columns. Estimations were obtained from 100,000 simulations per likelihood.

**Table S8.** Statistics of population scaled recombination rate ($\rho$, $bp^{-1}$) (mean ± standard error) over 10 Kbp non-overlapping windows along 19 chromosomes for the four *Populus* species

| Chromosome | *P. tremula* | *P. davidiana* | *P. tremuloides* | *P. trichocarpa* |
|---|---|---|---|---|
| Chr01 | 0.0129±0.0005 | 0.0095±0.0008 | 0.0264±0.0007 | 0.0106±0.0012 |
| Chr02 | 0.0137±0.0008 | 0.0097±0.0005 | 0.0234±0.0007 | 0.0094±0.0013 |
| Chr03 | 0.0139±0.0009 | 0.0103±0.0006 | 0.0299±0.0011 | 0.0090±0.0011 |
| Chr04 | 0.0151±0.0010 | 0.0075±0.0004 | 0.0255±0.0007 | 0.0097±0.0010 |
| Chr05 | 0.0123±0.0008 | 0.0087±0.0006 | 0.0246±0.0009 | 0.0090±0.0010 |
| Chr06 | 0.0123±0.0007 | 0.0088±0.0006 | 0.0256±0.0010 | 0.0106±0.0013 |
| Chr07 | 0.0144±0.0011 | 0.0107±0.0009 | 0.0284±0.0012 | 0.0104±0.0016 |
| Chr08 | 0.0133±0.0010 | 0.0080±0.0006 | 0.0262±0.0010 | 0.0113±0.0011 |
| Chr09 | 0.0139±0.0009 | 0.0083±0.0007 | 0.0281±0.0012 | 0.0091±0.0010 |
| Chr10 | 0.0128±0.0008 | 0.0094±0.0008 | 0.0249±0.0008 | 0.0098±0.0010 |
| Chr11 | 0.0147±0.0010 | 0.0103±0.0010 | 0.0305±0.0011 | 0.0090±0.0011 |
| Chr12 | 0.0162±0.0014 | 0.0093±0.0009 | 0.0323±0.0017 | 0.0119±0.0018 |
| Chr13 | 0.0144±0.0009 | 0.0107±0.0007 | 0.0291±0.0013 | 0.0105±0.0012 |
| Chr14 | 0.0118±0.0007 | 0.0091±0.0008 | 0.0294±0.0010 | 0.0111±0.0014 |
| Chr15 | 0.0154±0.0011 | 0.0111±0.0015 | 0.0283±0.0010 | 0.0093±0.0009 |
| Chr16 | 0.0183±0.0030 | 0.0142±0.0014 | 0.0312±0.0017 | 0.0131±0.0020 |
| Chr17 | 0.0158±0.0009 | 0.0102±0.0007 | 0.0309±0.0015 | 0.0107±0.0029 |
| Chr18 | 0.0139±0.0010 | 0.0097±0.0011 | 0.0273±0.0012 | 0.0118±0.0016 |
| Chr19 | 0.0156±0.0012 | 0.0116±0.0008 | 0.0312±0.0015 | 0.0067±0.0010 |
| Genomewide | 0.0139±0.0002 | 0.0096±0.0002 | 0.0273±0.0002 | 0.0102±0.0003 |

**Table S9.** Statistics of $F_{ST}$ (mean ± standard error) over 10 Kbp non-overlapping windows along 19 chromosomes between pairs of the four *Populus* species

| **Chromosome** | **P.tra-P.dav** | **P.tra-P.trs** | **P.dav-P.trs** | **P.tra-P.tri** | **P.dav-P.tri** | **P.trs-P.tri** |
|---|---|---|---|---|---|---|
| Chr01 | 0.3107±0.0022 | 0.3776±0.0024 | 0.3693±0.0023 | 0.7652±0.0013 | 0.7538±0.0013 | 0.7473±0.0013 |
| Chr02 | 0.3381±0.0030 | 0.3953±0.0031 | 0.3855±0.0030 | 0.7837±0.0015 | 0.7691±0.0015 | 0.7655±0.0015 |
| Chr03 | 0.3056±0.0036 | 0.3786±0.0034 | 0.3652±0.0034 | 0.7621±0.0021 | 0.7507±0.0021 | 0.7442±0.0020 |
| Chr04 | 0.3183±0.0034 | 0.3789±0.0036 | 0.3730±0.0035 | 0.7642±0.0020 | 0.7577±0.0020 | 0.7423±0.0020 |
| Chr05 | 0.3312±0.0033 | 0.3926±0.0032 | 0.3884±0.0031 | 0.7770±0.0018 | 0.7680±0.0018 | 0.7576±0.0018 |
| Chr06 | 0.3164±0.0030 | 0.3879±0.0030 | 0.3750±0.0030 | 0.7768±0.0016 | 0.7639±0.0016 | 0.7592±0.0016 |
| Chr07 | 0.3324±0.0041 | 0.3922±0.0045 | 0.3825±0.0044 | 0.7671±0.0026 | 0.7601±0.0025 | 0.7512±0.0025 |
| Chr08 | 0.3414±0.0033 | 0.4072±0.0034 | 0.3997±0.0033 | 0.7839±0.0017 | 0.7713±0.0018 | 0.7629±0.0017 |
| Chr09 | 0.3360±0.0040 | 0.3849±0.0039 | 0.3760±0.0038 | 0.7838±0.0021 | 0.7726±0.0021 | 0.7591±0.0020 |
| Chr10 | 0.3330±0.0031 | 0.3942±0.0031 | 0.3762±0.0031 | 0.7832±0.0017 | 0.7702±0.0017 | 0.7616±0.0016 |
| Chr11 | 0.2839±0.0039 | 0.3634±0.0043 | 0.3526±0.0038 | 0.7523±0.0024 | 0.7421±0.0024 | 0.7327±0.0023 |
| Chr12 | 0.2966±0.0041 | 0.3665±0.0043 | 0.3585±0.0042 | 0.7585±0.0026 | 0.7498±0.0026 | 0.7374±0.0026 |
| Chr13 | 0.3023±0.0040 | 0.4030±0.0046 | 0.3831±0.0044 | 0.7572±0.0027 | 0.7438±0.0027 | 0.7439±0.0027 |
| Chr14 | 0.3423±0.0037 | 0.3864±0.0038 | 0.3802±0.0037 | 0.7713±0.0024 | 0.7646±0.0024 | 0.7506±0.0024 |
| Chr15 | 0.2991±0.0042 | 0.3677±0.0044 | 0.3676±0.0043 | 0.7460±0.0029 | 0.7396±0.0029 | 0.7304±0.0028 |
| Chr16 | 0.2742±0.0040 | 0.3695±0.0044 | 0.3577±0.0043 | 0.7516±0.0026 | 0.7400±0.0026 | 0.7346±0.0026 |
| Chr17 | 0.2504±0.0043 | 0.3674±0.0049 | 0.3475±0.0045 | 0.7354±0.0032 | 0.7263±0.0031 | 0.7206±0.0030 |
| Chr18 | 0.2755±0.0039 | 0.3735±0.0043 | 0.3516±0.0040 | 0.7574±0.0027 | 0.7431±0.0027 | 0.7397±0.0028 |
| Chr19 | 0.2333±0.0044 | 0.3386±0.0049 | 0.3096±0.0046 | 0.7233±0.0035 | 0.7085±0.0036 | 0.7003±0.0036 |
| Genomewide | 0.3125±0.0008 | 0.3828±0.0008 | 0.3718±0.0008 | 0.7668±0.0005 | 0.7559±0.0005 | 0.7479±0.0005 |

Note: P. tra-*P. tremula*; P. dav-*P. davidiana*; P. trs-*P. tremuloides*; P. tri-*P. trichocarpa*.

**Table S10.** Statistics of $d_{xy}$ (mean ± standard error) over 10 Kbp non-overlapping windows along 19 chromosomes between pairs of the four *Populus* species

| Chromosome | P.tra-P.dav | P.tra-P.trs | P.dav-P.trs | P.tra-P.tri | P.dav-P.tri | P.trs-P.tri |
|---|---|---|---|---|---|---|
| Chr01 | 0.0213±0.0002 | 0.0244±0.0002 | 0.0246±0.0002 | 0.0440±0.0003 | 0.0436±0.0003 | 0.0435±0.0002 |
| Chr02 | 0.0206±0.0002 | 0.0232±0.0002 | 0.0237±0.0002 | 0.0427±0.0003 | 0.0423±0.0003 | 0.0422±0.0003 |
| Chr03 | 0.0197±0.0002 | 0.0222±0.0002 | 0.0226±0.0002 | 0.0413±0.0003 | 0.0410±0.0003 | 0.0409±0.0003 |
| Chr04 | 0.0199±0.0002 | 0.0225±0.0002 | 0.0228±0.0002 | 0.0413±0.0003 | 0.0410±0.0003 | 0.0410±0.0003 |
| Chr05 | 0.0201±0.0002 | 0.0228±0.0002 | 0.0231±0.0002 | 0.0420±0.0003 | 0.0417±0.0003 | 0.0416±0.0003 |
| Chr06 | 0.0205±0.0002 | 0.0230±0.0002 | 0.0235±0.0002 | 0.0422±0.0003 | 0.0418±0.0003 | 0.0417±0.0003 |
| Chr07 | 0.0186±0.0002 | 0.0210±0.0002 | 0.0213±0.0002 | 0.0397±0.0003 | 0.0393±0.0004 | 0.0393±0.0003 |
| Chr08 | 0.0200±0.0002 | 0.0227±0.0002 | 0.0231±0.0003 | 0.0418±0.0003 | 0.0415±0.0003 | 0.0414±0.0003 |
| Chr09 | 0.0187±0.0002 | 0.0211±0.0002 | 0.0213±0.0002 | 0.0397±0.0003 | 0.0394±0.0003 | 0.0394±0.0003 |
| Chr10 | 0.0201±0.0002 | 0.0227±0.0002 | 0.0231±0.0002 | 0.0421±0.0003 | 0.0417±0.0003 | 0.0417±0.0003 |
| Chr11 | 0.0185±0.0002 | 0.0209±0.0002 | 0.0212±0.0003 | 0.0394±0.0004 | 0.0391±0.0004 | 0.0391±0.0004 |
| Chr12 | 0.0186±0.0002 | 0.0209±0.0002 | 0.0212±0.0002 | 0.0395±0.0004 | 0.0392±0.0004 | 0.0392±0.0004 |
| Chr13 | 0.0187±0.0002 | 0.0211±0.0002 | 0.0214±0.0002 | 0.0397±0.0004 | 0.0393±0.0004 | 0.0393±0.0004 |
| Chr14 | 0.0189±0.0002 | 0.0214±0.0002 | 0.0217±0.0002 | 0.0401±0.0003 | 0.0398±0.0003 | 0.0398±0.0003 |
| Chr15 | 0.0187±0.0002 | 0.0211±0.0002 | 0.0214±0.0002 | 0.0398±0.0004 | 0.0394±0.0004 | 0.0394±0.0003 |
| Chr16 | 0.0186±0.0002 | 0.0209±0.0003 | 0.0212±0.0003 | 0.0393±0.0004 | 0.0390±0.0004 | 0.0390±0.0004 |
| Chr17 | 0.0183±0.0003 | 0.0208±0.0003 | 0.0211±0.0003 | 0.0390±0.0004 | 0.0387±0.0004 | 0.0387±0.0004 |
| Chr18 | 0.0185±0.0002 | 0.0208±0.0002 | 0.0211±0.0003 | 0.0394±0.0004 | 0.0391±0.0004 | 0.0391±0.0004 |
| Chr19 | 0.0180±0.0003 | 0.0207±0.0003 | 0.0209±0.0003 | 0.0386±0.0004 | 0.0382±0.0004 | 0.0383±0.0004 |
| Genomewide | 0.0197±0.0001 | 0.0222±0.0001 | 0.0226±0.0001 | 0.0412±0.0001 | 0.0409±0.0001 | 0.0408±0.0001 |

Note: P. tra-*P. tremula*; P. dav-*P. davidiana*; P. trs-*P. tremuloides*; P. tri-*P. trichocarpa*.

**Table S11.** Distribution of Spearman's rank correlation estimates (corr) of central summary statistics over 100 Kbp non-overlapping windows supporting linked selection shared among species as a central element shaping heterogeneous genomic diversity and differentiation. Subscripts i, j symbolize all possible combinations between two species i=1,…,n and j =i+1,…,n for within-species measures; Capital letters I, J symbolize inter-species statistics. Correlations were conducted between of all possible species comparisons I and J excluding comparisons with pseudo-replicated species (e.g. I=P.tra, P.dav; J= P.trs, P.tri). The measures of this table correspond to Figure 2B.

| Summary statistic | Mean | Median | Min | Max | Nr. Significance ($P$<0.001) | Nr. comparisons |
|---|---|---|---|---|---|---|
| Between-species comparison | | | | | | |
| corr($\pi_i$, $\pi_j$) | 0.7051 | 0.7063 | 0.5858 | 0.8039 | 6 | 6 |
| corr($\rho_i$, $\rho_j$) | 0.1801 | 0.1716 | 0.1111 | 0.2619 | 6 | 6 |
| corr($\mu$, $\pi_i$) | -0.0304 | -0.0297 | -0.0432 | -0.0192 | 0 | 4 |
| corr($\mu$, $\rho_i$) | 0.0236 | 0.0254 | 0.0061 | 0.0377 | 0 | 4 |
| Inter-species divergence comparison | | | | | | |
| corr($F_{ST\ I}$, $F_{ST\ J}$) | 0.5806 | 0.5592 | 0.3942 | 0.8304 | 15 | 15 |
| corr($d_{xy\ I}$, $d_{xy\ J}$) | 0.7650 | 0.6756 | 0.6321 | 0.9952 | 15 | 15 |
| corr($\pi_{i\ or\ j}$, $F_{ST\ I=i,j}$) | -0.5753 | -0.5474 | -0.7518 | -0.4275 | 12 | 12 |
| corr($\pi_{i\ or\ j}$, $d_{xy\ I=i,j}$) | 0.0502 | 0.0565 | 0.0120 | 0.0853 | 5 | 12 |
| corr($F_{ST\ I}$, $d_{xy\ I}$) | 0.0652 | 0.0668 | 0.0312 | 0.0806 | 4 | 6 |

Note: $\pi$ - Nucleotide diversity; $\rho$ - population-scaled recombination rate; $\mu$ – mutation rate by calculating the number of fixed differences per 'neutral' site (four-fold synonymous sites) between *P. tremula* and *P. trichocarpa*; $F_{ST}$ – interspecies genetic divergence; $d_{xy}$ – pairwise nucleotide divergence between species.

**Table S12.** Average topology weightings for each of the three topologies over 10 and 100 Kbp non-overlapping windows along 19 chromosomes.

| | **Topo1** | | **Topo2** | | **Topo3** | |
|---|---|---|---|---|---|---|
| | 10 Kbp | 100 Kbp | 10 Kbp | 100 Kbp | 10 Kbp | 100 Kbp |
| Chr01 | 0.2769 | 0.1966 | 0.5422 | 0.7628 | 0.1808 | 0.0404 |
| Chr02 | 0.3067 | 0.2481 | 0.5231 | 0.6955 | 0.1701 | 0.0563 |
| Chr03 | 0.2701 | 0.1807 | 0.5496 | 0.7433 | 0.1802 | 0.0758 |
| Chr04 | 0.2633 | 0.1503 | 0.5499 | 0.7892 | 0.1867 | 0.0604 |
| Chr05 | 0.2952 | 0.1867 | 0.5275 | 0.7520 | 0.1772 | 0.0612 |
| Chr06 | 0.2740 | 0.1905 | 0.5276 | 0.7675 | 0.1983 | 0.0418 |
| Chr07 | 0.2644 | 0.1259 | 0.5470 | 0.8004 | 0.1885 | 0.0736 |
| Chr08 | 0.2773 | 0.1562 | 0.5600 | 0.8147 | 0.1626 | 0.0289 |
| Chr09 | 0.2933 | 0.1722 | 0.5303 | 0.8070 | 0.1763 | 0.0207 |
| Chr10 | 0.2638 | 0.1625 | 0.5487 | 0.7651 | 0.1874 | 0.0722 |
| Chr11 | 0.2636 | 0.1598 | 0.5566 | 0.7462 | 0.1796 | 0.0939 |
| Chr12 | 0.2620 | 0.2154 | 0.5431 | 0.7215 | 0.1948 | 0.0629 |
| Chr13 | 0.2544 | 0.1562 | 0.5623 | 0.7581 | 0.1831 | 0.0855 |
| Chr14 | 0.2586 | 0.1977 | 0.5601 | 0.7496 | 0.1813 | 0.0526 |
| Chr15 | 0.2672 | 0.1907 | 0.5620 | 0.7589 | 0.1706 | 0.0503 |
| Chr16 | 0.2601 | 0.1905 | 0.5532 | 0.7659 | 0.1866 | 0.0434 |
| Chr17 | 0.2064 | 0.0946 | 0.6035 | 0.7908 | 0.1899 | 0.1145 |
| Chr18 | 0.2519 | 0.1440 | 0.5509 | 0.7888 | 0.1971 | 0.0671 |
| Chr19 | 0.2256 | 0.1118 | 0.5764 | 0.8091 | 0.1979 | 0.0789 |
| All | 0.2700 | 0.1757 | 0.5471 | 0.7648 | 0.1827 | 0.0594 |

Topo1: (((*P. tremula*, *P. tremuloides*), *P. davidiana*), *P. trichocarpa*)

Topo2: (((*P. tremula*, *P. davidiana*), *P. tremuloides*), *P. trichocarpa*)

Topo3: (((*P. davidiana*, *P. tremuloides*), *P. tremula*), *P. trichocarpa*)

**Table S13.** The number and percentage of 10 Kbp and 100 Kbp non-overlapping windows with a complete monophyly (weighting of 1) for each of the three topology, or with an unresolved tree.

| Windows | Top1 | | Top2 | | Top3 | | Unresolved | |
|---|---|---|---|---|---|---|---|---|
| | count | % all trees | count | % all trees | count | % all trees | count | % all trees |
| 10 Kbp | 1722 | 6.34% | 7776 | 28.62% | 603 | 2.22% | 17071 | 62.83% |
| 100 Kbp | 417 | 12.19% | 2403 | 70.22% | 69 | 2.02% | 533 | 15.58% |

**Table S14.** Summary of *D*-statistics and $f_d$-statistics based (GATK) or not based (ANGSD) on called genotypes over 10 Kbp and 100 Kbp non-overlapping windows across the chromosomes.

| | ANGSD | | | GATK | |
|---|---|---|---|---|---|
| **10 Kbp** | ***D*** | **Z-score** | ***P*-value** | ***D*** | $f_d$ |
| Chr01 | 0.091994 | 14.004665 | 0.000000 | 0.1118 | 0.0382 |
| Chr02 | 0.120093 | 12.296052 | 0.000000 | 0.1609 | 0.0498 |
| Chr03 | 0.068018 | 6.197107 | 0.000000 | 0.0964 | 0.0289 |
| Chr04 | 0.072098 | 7.777966 | 0.000000 | 0.0883 | 0.0292 |
| Chr05 | 0.091766 | 9.531925 | 0.000000 | 0.1160 | 0.0386 |
| Chr06 | 0.092296 | 10.352621 | 0.000000 | 0.1085 | 0.0352 |
| Chr07 | 0.055489 | 4.687780 | 0.000003 | 0.0905 | 0.0255 |
| Chr08 | 0.121079 | 12.037246 | 0.000000 | 0.1305 | 0.0417 |
| Chr09 | 0.094563 | 7.747383 | 0.000000 | 0.1202 | 0.0354 |
| Chr10 | 0.073900 | 7.588940 | 0.000000 | 0.0987 | 0.0302 |
| Chr11 | 0.078321 | 7.327218 | 0.000000 | 0.1073 | 0.0375 |
| Chr12 | 0.073825 | 6.565818 | 0.000000 | 0.1049 | 0.0361 |
| Chr13 | 0.061724 | 5.081574 | 0.000000 | 0.0842 | 0.0245 |
| Chr14 | 0.064805 | 5.822742 | 0.000000 | 0.0976 | 0.0286 |
| Chr15 | 0.077465 | 6.456604 | 0.000000 | 0.1020 | 0.0350 |
| Chr16 | 0.093687 | 7.988304 | 0.000000 | 0.1156 | 0.0359 |
| Chr17 | 0.012130 | 1.099662 | 0.271479 | 0.0389 | 0.0168 |
| Chr18 | 0.062434 | 5.573972 | 0.000000 | 0.0939 | 0.0308 |
| Chr19 | 0.019555 | 1.892397 | 0.058438 | 0.0542 | 0.0198 |
| All | 0.080783 | 34.075402 | 0.000000 | 0.1061 | 0.0341 |
| | **ANGSD** | | | **GATK** | |
| **100 Kbp** | ***D*** | **Z-score** | ***P*-value** | ***D*** | $f_d$ |
| Chr01 | 0.091979 | 11.259731 | 0.000000 | 0.1260 | 0.0367 |
| Chr02 | 0.120091 | 10.019572 | 0.000000 | 0.1756 | 0.0485 |
| Chr03 | 0.067991 | 5.038807 | 0.000000 | 0.0930 | 0.0283 |
| Chr04 | 0.072148 | 6.317156 | 0.000000 | 0.0871 | 0.0260 |
| Chr05 | 0.091829 | 6.819855 | 0.000000 | 0.1350 | 0.0385 |
| Chr06 | 0.092298 | 8.593502 | 0.000000 | 0.1393 | 0.0374 |
| Chr07 | 0.055524 | 3.399395 | 0.000675 | 0.0847 | 0.0208 |
| Chr08 | 0.121031 | 9.354301 | 0.000000 | 0.1616 | 0.0423 |
| Chr09 | 0.094541 | 7.218234 | 0.000000 | 0.1348 | 0.0368 |
| Chr10 | 0.073884 | 6.313919 | 0.000000 | 0.0907 | 0.0246 |
| Chr11 | 0.078615 | 6.099255 | 0.000000 | 0.0906 | 0.0267 |
| Chr12 | 0.073756 | 4.773741 | 0.000002 | 0.0948 | 0.0292 |
| Chr13 | 0.061579 | 3.914238 | 0.000091 | 0.0925 | 0.0244 |
| Chr14 | 0.064734 | 4.506067 | 0.000007 | 0.0883 | 0.0238 |
| Chr15 | 0.077518 | 4.445973 | 0.000009 | 0.1213 | 0.0353 |
| Chr16 | 0.093661 | 5.181017 | 0.000000 | 0.1128 | 0.0354 |

| | | | | | |
|---|---|---|---|---|---|
| Chr17 | 0.012161 | 0.774549 | 0.438606 | 0.0375 | 0.0110 |
| Chr18 | 0.062472 | 4.536191 | 0.000006 | 0.0977 | 0.0287 |
| Chr19 | 0.019588 | 1.565904 | 0.117371 | 0.0397 | 0.0155 |
| All | 0.080787 | 26.455704 | 0.000000 | 0.111703 | 0.0314 |

**Table S15.** Relationship between nucleotide diversity and coding density. Spearman's correlations between nucleotide diversity and coding density at two different scales (10 Kbp and 100 Kbp) in four *Populus* species. Also shown are partial correlation results after controlling for recombination rate (used the recombination of *P. tremula* as representative) and GC content.

| Species | Spearman's correlation between diversity and coding density | | Spearman's partial correlation controlling for recombination rate and GC content | |
|---|---|---|---|---|
| | 10 Kbp | 100 Kbp | 10 Kbp | 100 Kbp |
| *P. tremula* | -0.3437*** | -0.3637*** | -0.1773*** | -0.2439*** |
| *P. davidiana* | -0.3329*** | -0.3016*** | -0.1944*** | -0.2016*** |
| *P. tremuloides* | -0.3678*** | -0.3621*** | -0.2046*** | -0.2615*** |
| *P. trichocarpa* | -0.3705*** | -0.3240*** | -0.1363*** | -0.1647*** |

*$P$<0.01

**$P$<0.001

***$P$<0.0001

**Table S16.** Relationship between nucleotide diversity and recombination rate. Spearman's correlations between nucleotide diversity and recombination rate (used the recombination of *P. tremula* as representative) at two different scales (10 Kbp and 100 Kbp) in four *Populus* species. Also shown are partial correlation results after cntrolling for coding density and GC content.

| Species | Spearman's correlation between diversity and recombination rate | | Spearman's partial correlation controlling for coding density and GC content | |
|---|---|---|---|---|
| | 10 Kbp | 100 Kbp | 10 Kbp | 100 Kbp |
| *P. tremula* | 0.3024*** | 0.1928*** | 0.3231*** | 0.2468*** |
| *P. davidiana* | 0.3377*** | 0.2386*** | 0.3825*** | 0.2998*** |
| *P. tremuloides* | 0.3217*** | 0.1955*** | 0.3471*** | 0.2684*** |
| *P. trichocarpa* | 0.1439*** | 0.0735*** | 0.0981*** | 0.1258*** |

**P*<0.01

***P*<0.001

****P*<0.0001

**Table S17.** Relationship between genetic divergence ($F_{ST}$) and coding density. Spearman's correlations between $F_{ST}$ and coding density at two different scales (10 Kbp and 100 Kbp) in four *Populus* species. Also shown are partial correlation results after controlling for recombination rate (used the recombination of *P. tremula* as representative)and GC content.

| Species pairs | Spearman's correlation between $F_{ST}$ and coding density | | Spearman's partial correlation controlling for recombination rate and GC content | |
|---|---|---|---|---|
| | 10 Kbp | 100 Kbp | 10 Kbp | 100 Kbp |
| *P. tremula* vs. *P. davidiana* | -0.043*** | -0.0881*** | -0.0591*** | -0.0929*** |
| *P. tremula* vs. *P. tremuloides* | 0.0069 | -0.0375 | -0.0008 | -0.0296 |
| *P. davidiana* vs. *P. tremuloides* | -0.0174 | -0.1075*** | -0.0107 | -0.0994*** |
| *P. tremula* vs. *P. trichocarpa* | 0.0191* | -0.0574* | -0.0419*** | -0.1185*** |
| *P. davidiana* vs. *P. trichocarpa* | -0.0085 | -0.1302*** | -0.0488*** | -0.1823*** |
| *P. tremuloides* vs. *P. trichocarpa* | 0.0077 | -0.1102*** | -0.0390*** | -0.1582*** |

*$P$<0.01
**$P$<0.001
***$P$<0.0001

**Table S18.** Relationship between genetic divergence ($F_{ST}$) and recombination rate. Spearman's correlations between $F_{ST}$ and recombination rate (used the recombination of *P. tremula* as representative) at two different scales (10 Kbp and 100 Kbp) in four *Populus* species. Also shown are partial correlation results after cntrolling for coding density and GC content.

| Species pairs | Spearman's correlation between $F_{ST}$ and recombination rate | | Spearman's partial correlation controlling for coding density and GC content | |
|---|---|---|---|---|
| | 10 Kbp | 100 Kbp | 10 Kbp | 100 Kbp |
| *P. tremula* vs. *P. davidiana* | -0.1631*** | -0.1671*** | -0.1639*** | -0.1492*** |
| *P. tremula* vs. *P. tremuloides* | -0.1743*** | -0.1633*** | -0.1743*** | -0.1556*** |
| *P. davidiana* vs. *P. tremuloides* | -0.1228*** | -0.1383*** | -0.1231*** | -0.1199*** |
| *P. tremula* vs. *P. trichocarpa* | -0.2073*** | -0.2181*** | -0.2081*** | -0.1967*** |
| *P. davidiana* vs. *P. trichocarpa* | -0.1496*** | -0.1874*** | -0.1501*** | -0.1560*** |
| *P. tremuloides* vs. *P.trichocarpa* | -0.1434*** | -0.1791*** | -0.1438*** | -0.1511*** |

*$P$<0.01

**$P$<0.001

***$P$<0.0001

**Table S19.** Relationship between genetic divergence ($d_{xy}$) and coding density. Spearman's correlations between $d_{xy}$ and coding density at two different scales (10 Kbp and 100 Kbp) in four *Populus* species. Also shown are partial correlation results after controlling for recombination rate (used the recombination of *P. tremula* as representative) and GC content.

| Species pairs | Spearman's correlation between $d_{xy}$ and coding density | | Spearman's partial correlation controlling for recombination rate and GC content | |
|---|---|---|---|---|
| | 10 Kbp | 100 Kbp | 10 Kbp | 100 Kbp |
| *P. tremula* vs. *P. davidiana* | $-0.0304^{***}$ | -0.0211 | $-0.0270^{**}$ | -0.0358 |
| *P. tremula* vs. *P. tremuloides* | $-0.0323^{***}$ | -0.0331 | $-0.0215^{*}$ | -0.0416 |
| *P. davidiana* vs. *P. tremuloides* | $-0.0346^{***}$ | -0.0423 | $-0.0265^{**}$ | $-0.0588^{*}$ |
| *P. tremula* vs. *P. trichocarpa* | $-0.0555^{***}$ | $-0.0866^{***}$ | $-0.0365^{***}$ | $-0.0893^{***}$ |
| *P. davidiana* vs. *P. trichocarpa* | $-0.0568^{***}$ | $-0.0874^{***}$ | $-0.0372^{***}$ | $-0.0895^{***}$ |
| *P. tremuloides* vs. *P. trichocarpa* | $-0.0557^{***}$ | $-0.0891^{***}$ | $-0.0371^{***}$ | $-0.0917^{***}$ |

$^{*}P<0.01$

$^{**}P<0.001$

$^{***}P<0.0001$

**Table S20.** Relationship between genetic divergence ($d_{xy}$) and recombination rate. Spearman's correlations between $d_{xy}$ and recombination rate (used the recombination of *P. tremula* as representative) at two different scales (10 Kbp and 100 Kbp) in four *Populus* species. Also shown are partial correlation results after cntrolling for coding density and GC content.

| Species pairs | Spearman's correlation between $d_{xy}$ and recombination rate | | Spearman's partial correlation controlling for coding density and GC content | |
|---|---|---|---|---|
| | 10 Kbp | 100 Kbp | 10 Kbp | 100 Kbp |
| *P. tremula* vs. *P. davidiana* | 0.0121 | -0.0194 | 0.0117 | -0.0126 |
| *P. tremula* vs. *P. tremuloides* | 0.0155 | -0.0207 | 0.0152 | -0.013 |
| *P. davidiana* vs. *P. tremuloides* | 0.0195* | 0.0258 | 0.0192* | -0.0148 |
| *P. tremula* vs. *P. trichocarpa* | 0.0118 | -0.0425 | 0.0113 | -0.0265 |
| *P. davidiana* vs. *P. trichocarpa* | 0.0122 | -0.0426 | 0.0116 | -0.0265 |
| *P. tremuloides* vs. *P. trichocarpa* | 0.0123 | -0.0415 | 0.0118 | -0.025 |

*$P$<0.01
**$P$<0.001
***$P$<0.0001

**Table S21.** Relationship between coding density and incomplete lineage sorting (ILS), topology weighting of species tree and admixture proportion ($f_d$). Spearman's correlations between these parameters and coding density at two different scales (10 Kbp and 100 Kbp) in four *Populus* species. Also shown are partial correlation results after controlling for recombination rate (used the recombination of *P. tremula* as representative) and GC content.

| | Spearman's correlation with coding density | | Spearman's partial correlation controlling for recombination rate and GC content | |
|---|---|---|---|---|
| | 10 Kbp | 100 Kbp | 10 Kbp | 100 Kbp |
| ILS | -0.0908*** | -0.1062*** | -0.0454*** | -0.0597* |
| Weighting of species tree | 0.0363*** | 0.1268*** | 0.0331*** | 0.1109*** |
| $f_d$ | -0.0747*** | -0.1588*** | -0.0456*** | -0.1540*** |

*$P$<0.01
**$P$<0.001
***$P$<0.0001

**Table S22.** Relationship between recombination rate and incomplete lineage sorting (ILS), topology weighting of species tree and admixture proportion ($f_d$). Spearman's correlations between these parameters and recombination rate (used the recombination of *P. tremula* as representative) at two different scales (10 Kbp and 100 Kbp) in four *Populus* species. Also shown are partial correlation results after controlling for coding density and GC content.

| | Spearman's correlation with recombination rate | | Spearman's partial correlation controlling for coding density and GC content | |
|---|---|---|---|---|
| | 10 Kbp | 100 Kbp | 10 Kbp | 100 Kbp |
| ILS | 0.1874*** | 0.0787*** | 0.1875*** | 0.0869*** |
| Weighting of species tree | 0.0315*** | 0.0755*** | 0.0318*** | 0.0559* |
| $f_d$ | -0.0122 | 0.0076 | -0.0125 | 0.0309 |

*$P$<0.01
**$P$<0.001
***$P$<0.0001

**Table S23.** Enriched Gene Ontology (GO) categories for genes located in windows identified by SweepFinder2 as being under positive selection in the three aspen species: *P. tremula*, *P. davidiana* and *P. tremuloides.*

| GO.ID | Term | Annotated | Significant | Expected | Fisher.p |
|---|---|---|---|---|---|
| GO:0006259 | DNA metabolic process | 332 | 13 | 4.34 | 0.00044 |
| GO:0090304 | nucleic acid metabolic process | 2366 | 49 | 30.96 | 0.00054 |
| GO:0044260 | cellular macromolecule metabolic process | 5705 | 97 | 74.65 | 0.00068 |
| GO:0006996 | organelle organization | 230 | 10 | 3.01 | 0.00091 |
| GO:0006725 | cellular aromatic compound metabolic process | 2939 | 56 | 38.46 | 0.00152 |
| GO:0046483 | heterocycle metabolic process | 2944 | 56 | 38.52 | 0.00159 |
| GO:0006400 | tRNA modification | 21 | 3 | 0.27 | 0.00247 |
| GO:0006777 | Mo-molybdopterin cofactor biosynthetic process | 6 | 2 | 0.08 | 0.00247 |
| GO:0019720 | Mo-molybdopterin cofactor metabolic process | 6 | 2 | 0.08 | 0.00247 |
| GO:0032324 | molybdopterin cofactor biosynthetic process | 6 | 2 | 0.08 | 0.00247 |
| GO:0042816 | vitamin B6 metabolic process | 6 | 2 | 0.08 | 0.00247 |
| GO:0042819 | vitamin B6 biosynthetic process | 6 | 2 | 0.08 | 0.00247 |
| GO:0043545 | molybdopterin cofactor metabolic process | 6 | 2 | 0.08 | 0.00247 |
| GO:0051189 | prosthetic group metabolic process | 6 | 2 | 0.08 | 0.00247 |
| GO:0006139 | nucleobase-containing compound metabolic process | 2815 | 53 | 36.83 | 0.00278 |
| GO:0044237 | cellular metabolic process | 7292 | 115 | 95.41 | 0.00292 |
| GO:1901360 | organic cyclic compound metabolic process | 3092 | 57 | 40.46 | 0.00297 |
| GO:0000902 | cell morphogenesis | 25 | 3 | 0.33 | 0.0041 |
| GO:0009653 | anatomical structure morphogenesis | 25 | 3 | 0.33 | 0.0041 |
| GO:0032989 | cellular component morphogenesis | 25 | 3 | 0.33 | 0.0041 |
| GO:0048869 | cellular developmental process | 25 | 3 | 0.33 | 0.0041 |
| GO:0016043 | cellular component organization | 437 | 13 | 5.72 | 0.00501 |
| GO:0006260 | DNA replication | 85 | 5 | 1.11 | 0.00513 |
| GO:0071840 | cellular component organization or biogenesis | 504 | 14 | 6.59 | 0.00658 |
| GO:0043170 | macromolecule metabolic process | 6307 | 100 | 82.52 | 0.00688 |
| GO:0051276 | chromosome organization | 134 | 6 | 1.75 | 0.00838 |
| GO:0006074 | (1->3)-beta-D-glucan metabolic process | 11 | 2 | 0.14 | 0.00867 |
| GO:0006075 | (1->3)-beta-D-glucan biosynthetic process | 11 | 2 | 0.14 | 0.00867 |
| GO:0006265 | DNA topological change | 11 | 2 | 0.14 | 0.00867 |
| GO:0040029 | regulation of gene expression, epigenetic | 11 | 2 | 0.14 | 0.00867 |
| GO:0009987 | cellular process | 9803 | 144 | 128.27 | 0.00927 |
